# *Ptr1* evolved convergently with *RPS2* and *Mr5* to mediate recognition of AvrRpt2 in diverse solanaceous species

**DOI:** 10.1101/2020.03.05.979484

**Authors:** Carolina Mazo-Molina, Samantha Mainiero, Benjamin J. Haefner, Ryland Bednarek, Jing Zhang, Ari Feder, Kai Shi, Susan R. Strickler, Gregory B. Martin

**Affiliations:** Boyce Thompson Institute for Plant Research, Ithaca, NY 14853, USA; Plant Pathology and Plant-Microbe Biology Section, School of Integrative Plant Science, Cornell University, Ithaca, NY 14853, USA; Department of Horticulture, Zhejiang University, Hangzhou 310058, China

**Author notes:** Corresponding author: G. B. Martin, Boyce Thompson Institute for Plant Research, Ithaca, NY 14853, USA. These authors made equal contributions and are considered co-first authors (Carolina Mazo-Molina, Samantha Mainiero).

**Keywords:** AvrRpt2, Disease resistance, Effector-triggered immunity, *Nicotiana benthamiana*, *Pseudomonas syringae*, *Ralstonia pseudosolanacearum*, Rin4, RipBN, *Solanum lycopersicoides*, *Solanum tuberosum*

## Abstract

The *Ptr1 (Pseudomonas tomato race 1)* locus in *Solanum lycopersicoides* confers resistance to strains of *Pseudomonas syringae* pv. tomato expressing AvrRpt2 and *Ralstonia pseudosolanacearum* expressing RipBN. Here we describe the identification and phylogenetic analysis of the *Ptr1* gene. A single recombinant among 585 F2 plants segregating for the *Ptr1* locus was discovered that narrowed the *Ptr1* candidates to eight nucleotide-binding leucine-rich repeat protein (NLR)-encoding genes. From analysis of the gene models in the *S. lycopersicoides* genome sequence and RNA-Seq data, two of the eight genes emerged as the strongest candidates for *Ptr1*. One of these two candidates was found to encode *Ptr1* based on its ability to mediate recognition of AvrRpt2 and RipBN when it was transiently expressed with these effectors in leaves of *Nicotiana glutinosa*. The ortholog of *Ptr1* in tomato and in *Solanum pennellii* is a pseudogene. However, a functional *Ptr1* ortholog exists in *N. benthamiana* and potato and both mediate recognition of AvrRpt2 and RipBN. In apple and Arabidopsis, recognition of AvrRpt2 is mediated by the Mr5 and RPS2 proteins, respectively. Phylogenetic analysis places *Ptr1* in a distinct clade compared to *Mr5* and *RPS2* and it therefore appears to have arisen by convergent evolution for recognition of AvrRpt2.

## Introduction

Bacterial speck disease of tomato, caused by *Pseudomonas syringae* pv. tomato *(Pst)*, is a persistent problem in production areas throughout the world where conditions are cool and moist (Jones, 1991, Pedley and Martin, 2003). The disease causes necrotic lesions (specks) on different parts of the plant including leaves, stems, flowers, and fruits. As a result, it can affect both fruit yield and quality, leading to significant economic losses (Jones, 1991). Genetic resistance to *Pst* is conferred by the *Pto* and *Prf* genes which encode a serine/threonine cytoplasmic kinase and a nucleotide-binding leucine-rich repeat (NLR) protein, respectively (Pedley and Martin, 2003). The Pto/Prf proteins form a complex that recognizes the type III effectors AvrPto or AvrPtoB expressed by race 0 *Pst* strains (Kim *et al*., 2002, Mucyn *et al*., 2009, Martin, 2012, Ntoukakis *et al*., 2013, Saur *et al*., 2015). The widespread use of the *Pto/Prf* genes since the 1980s appears to have exerted selection pressure on the pathogen and race 0 *Pst* strains are becoming less common. Instead, strains of race 1 *Pst*, which lack AvrPto and AvrPtoB, are becoming more prevalent and represent a new threat in tomato growing areas (Kunkeaw *et al*., 2010, Thapa and Coaker, 2016).

In the summer of 2015 a natural outbreak of bacterial speck disease occurred on a Cornell University research farm where 110 *Solanum lycopersicum* VF36 x *S. lycopersicoides* LA2951 introgression lines (ILs) were being grown (Canady *et al*., 2005, Kraus *et al*., 2017, Mazo-Molina *et al*., 2019). Two of the ILs, LA4245 and LA4277, remained free of the disease. These two lines each carry a large, partially overlapping, introgression on chromosome 4 from *S. lycopersicoides* and in later controlled inoculations were found to have resistance to race 1, but not race 0, strains of *Pst* (Canady *et al*., 2005, Mazo-Molina *et al*., 2019). The locus responsible was named *Ptr1* for *<u>P</u>. s. pv. tomato <u>r</u>ace <u>1</u>* resistance (Mazo-Molina *et al*., 2019). Subsequently, it was determined that the *Ptr1* locus mediated recognition of the effector AvrRpt2 which is present in all *Pst* race 1 strains for which a genome sequence is available (Mazo-Molina *et al*., 2019).

AvrRpt2 is a cysteine protease which is translocated by the *Pst* type III secretion system into host cells during the infection process (Mudgett and Staskawicz, 1999, Axtell *et al*., 2003). In Arabidopsis, AvrRpt2 degrades the *At*RIN4 protein, thereby activating RPS2-mediated defense responses (Axtell and Staskawicz, 2003, Mackey *et al*., 2003, Day *et al*., 2005). In apple, AvrRpt2 from *E. amylovora* is recognized by the resistance gene *Mr5*, although whether RIN4 is involved in this process is unknown (Fahrentrapp *et al*., 2012, Vogt *et al*., 2013). *RPS2* and *Mr5* have no sequence similarity and appear to be the result of convergent evolution (Fahrentrapp *et al*., 2012).

Tomato has three genes with similarity to *AtRIN4* that are expressed in leaves (*Sl*Rin4-1, *Sl*Rin4-2, and *Sl*Rin4-3) (Mazo-Molina *et al*., 2019). As in Arabidopsis, recognition of AvrRpt2 by the *Ptr1* locus was found to be mediated by the ability of AvrRpt2 to induce the degradation of tomato Rin4 proteins (Mazo-Molina *et al*., 2019). In addition, a mutagenesis analysis of AvrRpt2 revealed that, similar to RPS2, activation of Ptr1 required the cysteine proteolytic activity of AvrRpt2 (Mazo-Molina *et al*., 2019). Interestingly, the *Ptr1* locus was also able to mediate recognition of AvrRpt2 homologs present in a diverse array of different bacteria. In particular, the *Ptr1* locus recognized the effector RipBN in *Ralstonia pseuodsolanacearum* and conferred effective resistance to that vascular pathogen (Mazo-Molina *et al*., 2019).

A genome sequence for *Solanum lycopersicoides* LA2951 was generated and used to examine the introgression on chromosome 4 in LA4245 and LA4277 that carries the *Ptr1* locus (Mazo-Molina *et al*., 2019, Powell *et al*., 2020). A total of 16 genes which encode predicted NLR proteins were annotated in the introgression region shared between LA4245 and LA4277 and we hypothesized that one of these genes was *Ptr1* (Mazo-Molina *et al*., 2019). Here we present the identification of the *Ptr1* gene and show that it is not orthologous to either *RPS2* or *Mr5* and therefore appears to have arisen by convergent evolution to mediate recognition of AvrRpt2.

## Results

### Identifying candidates for the *Ptr1* gene

The introgression on chromosome 4 containing the *Ptr1* locus is maintained in heterozygous condition and we refer to such *Ptr1/ptr1* plants as LA4245-R and LA4277-R. Homozygous *Ptr1/Ptr1* plants are referred to as LA4245-Ro and LA4277-Ro and *ptr1/ptr1* plants are referred to as LA4245-S and LA4277-S (Mazo-Molina *et al*., 2019). The introgression region shared by LA4245-R and LA4277-R contains 16 annotated NLR-encoding genes in the current assembly of the *S. lycopersicoides* genome sequence (Powell *et al*., 2020). An additional ~50 NLR genes occur only in LA4277-R; the latter genes were not considered as candidates for *Ptr1* (**Figure 1A**). To identify candidates for the *Ptr1* gene, F2 progeny from a selfed LA4277-R plant were screened for recombinants. Plants were genotyped for the presence of the *S. lycopersicoides* introgression using DNA markers at 7.14, 9.94, and 69.13 megabases (Mb) (**Figure 1A**). From 585 F2 plants, one plant, LA4277-159, was identified with a recombination between markers 7.14 and 9.94. An additional marker at 9.15 Mb further delimited the region of the recombination and indicated this plant retained just a small portion of the *S. lycopersicoides* segment shared by both LA4245-R and LA4277-R (**Figure 1B**). This segment contains eight NLR-encoding genes.

**Figure 1.**
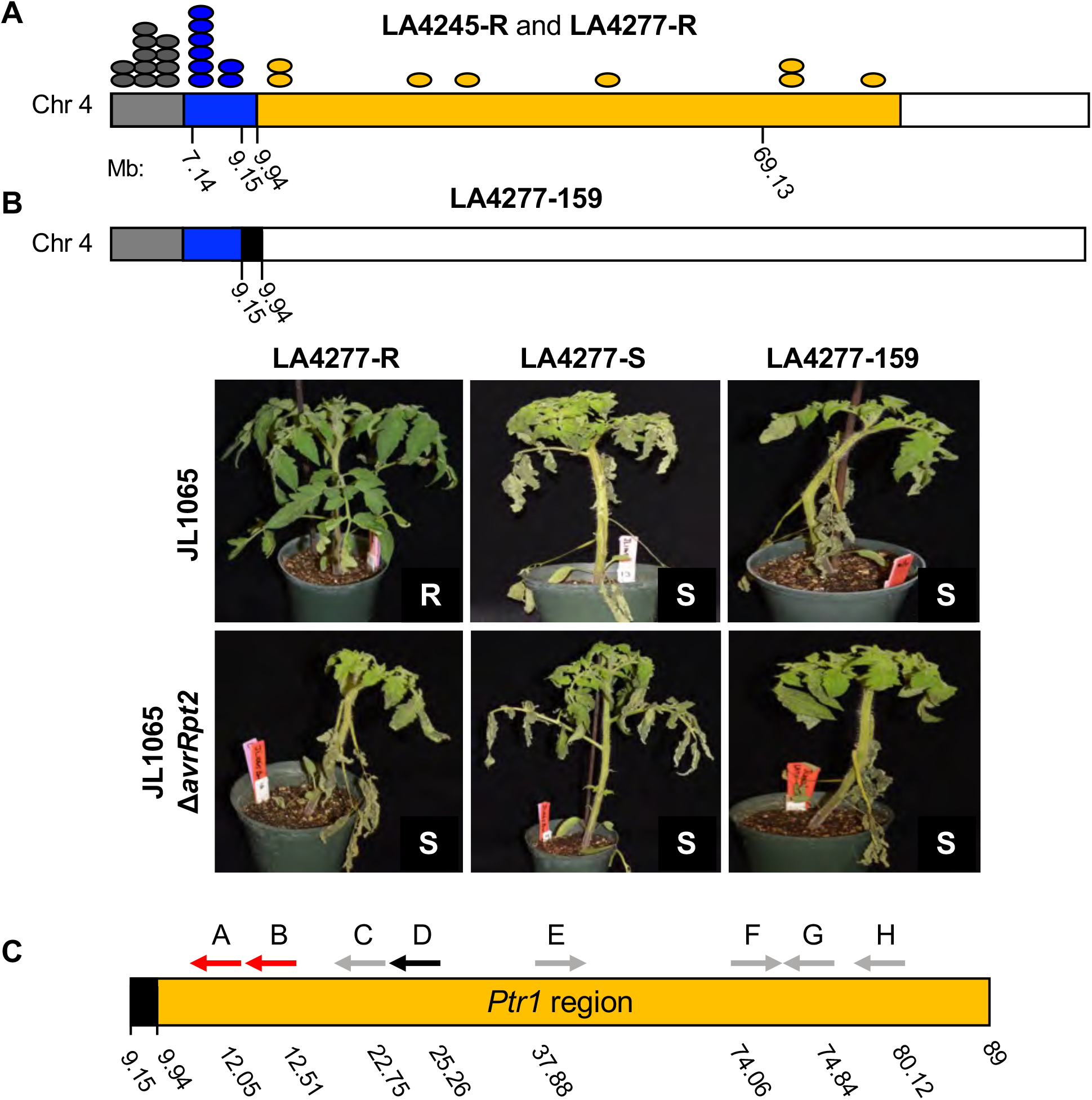
Delimiting the region containing *Ptr1* using progeny of selfed LA4277-R (*Ptr1/ptr1*) plants. **A)** Schematic of *S. lycopersicoides* introgression on chromosome 4 in LA4245-R (gold and blue regions) and LA4277-R (gold, blue, and gray regions). The region in white is derived from the tomato parent VF36. Distribution of NLR-encoding genes unique to LA4277-R (gray dots, representing ~50 NLR genes) and those shared with LA4245-R (gold and blue dots) is shown. Indicated are the locations of the DNA markers (megabases, Mb) used to identify recombinants. **B)** LA4277-159 derived from a selfed LA4277-R plant has a recombination event between markers at 9.15 and 9.94 Mb (black box). LA4277-R, LA4277-S, and LA4277-159 plants were vacuum infiltrated with *Pst* JL1065 or *Pst* JL1065△*avrRpt2*. Photos were taken 6 days after inoculation and are representative of three independent experiments. R, resistant; S, susceptible. **C)** The region defined as containing *Ptr1* contains eight NLR gene models, A-H. Their positions and orientations within the *Ptr1* region are shown with the coordinates (Mb) corresponding to the start codon of each gene model. Transcripts were not detectable in leaves from the genes depicted in gray and candidate D is a pseudogene.

To determine if LA4277-159 contained the *Ptr1* locus, selfed progeny of the recombinant plant that contain the *S. lycopersicoides* 0 – 9.94 Mb segment were identified by using DNA markers. These plants, and controls, were then vacuum infiltrated with *Pst* JL1065 (which has *avrRpt2*) and JL1065Δ*avrRpt2*, which lacks *avrRpt2*. As expected, LA4277-R, LA4277-S, and the LA4277-159 progeny all developed extensive disease upon inoculation with JL1065Δ*avrRpt2* (**Figure 1B**). However, when infiltrated with JL1065, LA4277-159 progeny and LA4277-S developed disease whereas LA4277-R plants remained disease-free. Considering the location of the recombination event in LA4277-159 this susceptible phenotype eliminated the eight NLR genes in the LA4277-159 *S. lycopersicoides* segment as candidates for *Ptr1* and indicated *Ptr1* is one of the eight NLR genes lying between marker 9.94 and the end of the introgression at 89 Mb (**Figure 1C**). These genes were named *A*, *B*, *C, D, E, F, G*, and *H* based on their sequential location on chromosome 4 (**Figure 1C** and **Table S1**).

We next used RNA-Seq analysis to determine which of the eight *Ptr1* candidates is expressed in leaves of a LA4277-Ro (*Ptr1/Ptr1*) plant (**Table S2**). Seven-week-old plants were vacuum infiltrated with JL1065 and the abundance of transcripts was determined by 3’ RNA-Seq. Transcripts of only three candidates, *A*, *B* and *D* were detectable by this method (**Table S2**). Each of these genes was PCR amplified from cDNA derived from a LA4277-Ro plant and sequenced to determine whether they were the same as annotated in the *S. lycopersicoides* LA2951 genome sequence. In addition, an LA4277 genome sequence was generated and assembled as a comparison. Candidate D was found to be a pseudogene as it contained multiple mutations disrupting the reading frame. The sequence of candidate A was identical between LA2951 and LA4277 whereas one SNP occurred in candidate B which changed an alanine in LA2951 to a valine in LA4277.

### Candidate A mediates recognition of AvrRpt2 and RipBN in *Nicotiana glutinosa*

Candidates A and B were cloned into a binary vector under control of the CaMV 35S promoter and tested by *Agrobacterium*-mediated transient transformation (‘agroinfiltration’). *Nicotiana glutinosa* was used in these experiments because it was reported previously that AvrRpt2 does not cause cell death in this species whereas it does in *N. benthamiana* and this would have interfered with the assays (Mudgett and Staskawicz, 1999, Day *et al*., 2005, Kessens *et al*., 2014).

Syringe infiltration into leaves of *N. glutinosa* of a relatively low titer (OD_600_ = 0.025) of *Agrobacterium* carrying constructs of candidate A or B expressed from the CaMV 35S promoter caused no observable host response (**Figure 2A**). However, co-expression of candidate A with either AvrRpt2 or RipBN induced cell death two days after infiltration. No cell death was observed when candidate A was co-expressed with the protease-inactive protein AvrRpt2(C122A). Candidate B did not induce cell death when co-expressed with any of the effector proteins (**Figure 2A**).

**Figure 2.**
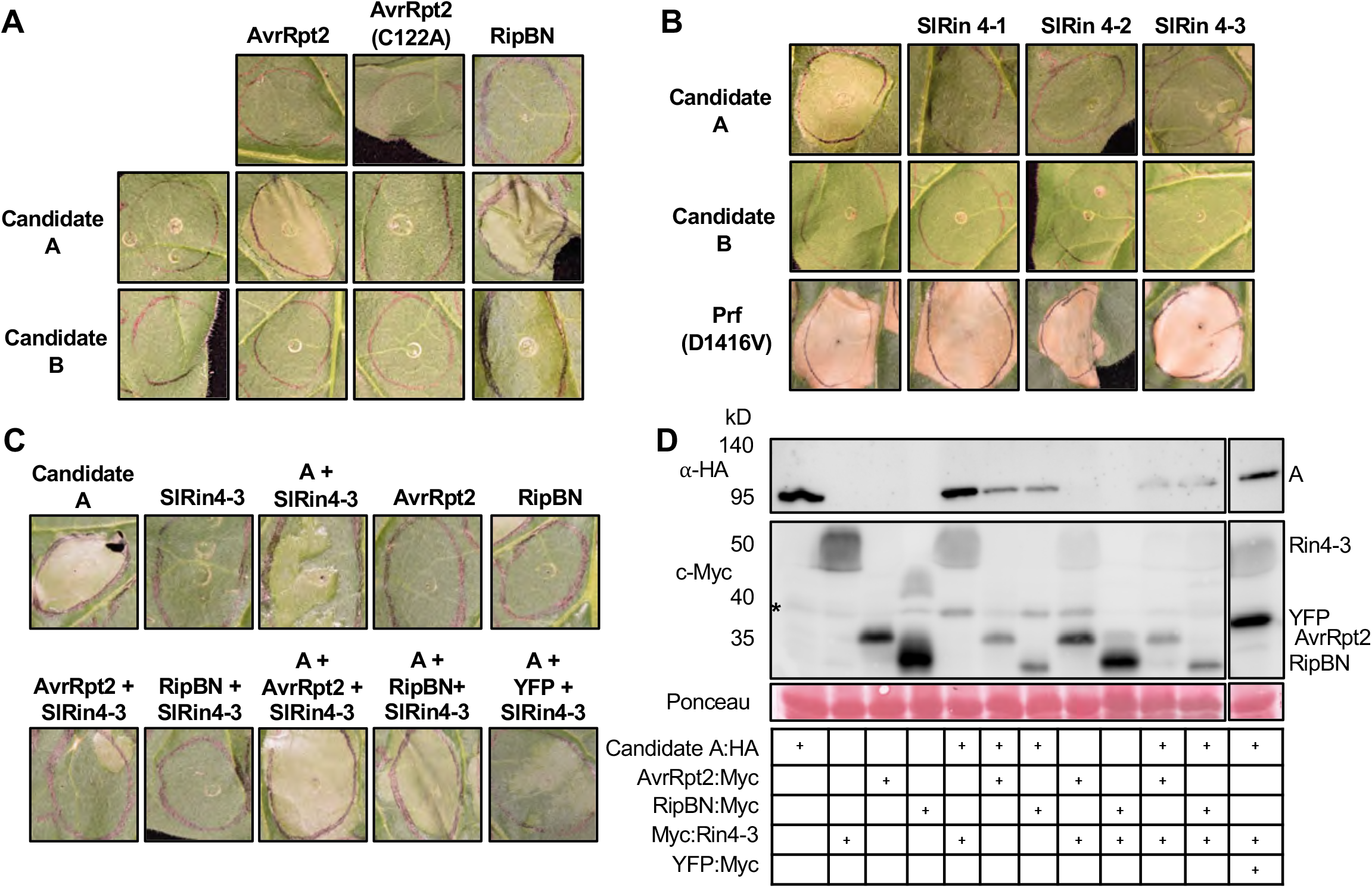
Candidate A mediates recognition of AvrRpt2 and RipBN in *Nicotiana glutinosa*. **A)** *N. glutinosa* leaves were syringe-infiltrated with *Agrobacterium* strains carrying either *A:HA* or *B:HA* (OD_600_ = 0.025), and *AvrRpt2:Myc, AvrRpt2(C122A):Myc* or *RipBN:Myc* (OD_600_ = 0.05). Photographs were taken 48 hr after infiltration. **B)** *N. glutinosa* leaves were syringe-infiltrated with *Agrobacterium* strains expressing *A:HA* or *B:HA* (OD_600_ = 0.1), *Prf(D1416V):HA* (OD_600_ = 0.1) or *Myc:*Sl*Rin4* (OD_600_ = 0.2). Photographs for candidates A and B were taken 48 hr after infiltration and 4 days after infiltration for Prf(D1416V). **C)** *N. glutinosa* leaves were syringe-infiltrated with *Agrobacterium* strains carrying either *A:HA* (OD_600_ = 0.1), *AvrRpt2:Myc* (OD_600_ = 0.05), *RipBN:Myc* (OD_600_ = 0.05), *YFP:Myc* (OD_600_ = 0.05) or *Myc*:*SlRin4-3* (OD_600_ = 0.2). Photographs were taken 48 hr after infiltration. **D)** Immunoblot analysis of protein extracts isolated from *N. benthamiana* leaves agroinfiltrated with *A:HA* (OD_600_ = 0.1), *SlRin4-3:Myc* (OD_600_ = 0.2), *AvrRpt2:Myc* (OD_600_ = 0.05), *RipBN:Myc* (OD_600_ = 0.05), or *YFP:Myc* (OD_600_ = 0.05). Samples were collected 28 hr after infiltration. Total proteins extracted from infiltrated leaves were subjected to immunoblotting using an ∝-HA antibody to detect candidate A and c-Myc antibody to detect SlRin4-3, AvrRpt2, RipBN, and YFP. +, construct was included in the experiment. Protein masses are indicated at the left of the blot. Ponceau staining shows amount of protein loaded in each lane. The asterisk indicates a nonspecific band across the immunoblot. All genes were expressed from the CaMV 35S promoter. Data shown are representative of three independent experiments, using three biological replicates and infiltrating two leaves per plant.

RPS2 can cause cell death on its own when overexpressed in *N. benthamiana* leaves and this cell death is suppressed by co-expression of *At*RIN4 (Day *et al*., 2005). We therefore syringe infiltrated *Agrobacterium* strains carrying the candidate A and B constructs at a four-fold higher titer (OD_600_ = 0.1). At this titer, candidate A, but not candidate B, induced strong cell death on its own. Tomato has three *Rin4* genes that are expressed in leaves (*SlRin4-1*, *SlRin4-2*, and *SlRin4-3*) (**Figure S1A;** (Mazo-Molina *et al*., 2019)). To test whether the tomato Rin4 proteins can suppress cell death caused by candidate A, *Sl*Rin4-1, *Sl*Rin4-2 and *Sl*Rin 4-3 were expressed with candidate A in *N. glutinosa* leaves. Each of these proteins completely suppressed candidate A-mediated cell death (**Figure 2B** and **Figure S1B**). Candidate B did not cause cell death even at higher *Agrobacterium* titers nor was it affected by co-expression of *Sl*Rin4-3 or either of the effectors (**Figure 2B** and **Figure S2**). A constitutive-active version of Prf was used as a positive control for cell death in these experiments and this response was not suppressed by co-expression with the tomato *Sl*Rin4 proteins (**Figure 2B**).

In Arabidopsis, AvrRpt2 causes degradation of *At*RIN4 resulting in the activation of RPS2 (Axtell and Staskawicz, 2003, Mackey *et al*., 2003). To test whether this is the case for candidate A, we co-expressed candidate A with *Sl*Rin4-3 with or without the effectors AvrRpt2 and RipBN (**Figure 2C**). Co-expression of either one of the effectors with candidate A and *Sl*Rin4-3 restored candidate A-induced cell death and this was correlated with a decrease in the abundance of Rin4 protein (**Figure 2D**). All proteins were found to be expressed as expected by immunoblotting (**Figure 2D**). These experiments demonstrate that candidate A is the *Ptr1* gene. The predicted Ptr1 protein is a coiled-coil NLR with 13 leucine-rich repeats and all of the other hallmark motifs of this class of protein (**Figure S3**).

### *Ptr1* is a pseudogene in tomato and *S. pennellii*, but intact *Ptr1* genes occur in other solanaceous species

No accessions of tomato or its close wild relatives are known that recognize AvrRpt2. We searched the genome sequences of the tomato reference genome (Heinz 1706) and its wild relative *S. pennellii* (LA0716; (Bolger *et al*., 2014a)) and found that, while both have a clear ortholog of *Ptr1* (with greater than 95% identical nucleotide sequences), a small deletion abolishes the *Ptr1* start codon in both species and multiple other nonsense and frameshift mutations disrupt the reading frames of these *Ptr1* genes (**Figure 3A** and **Figure S4**).

**Figure 3.**
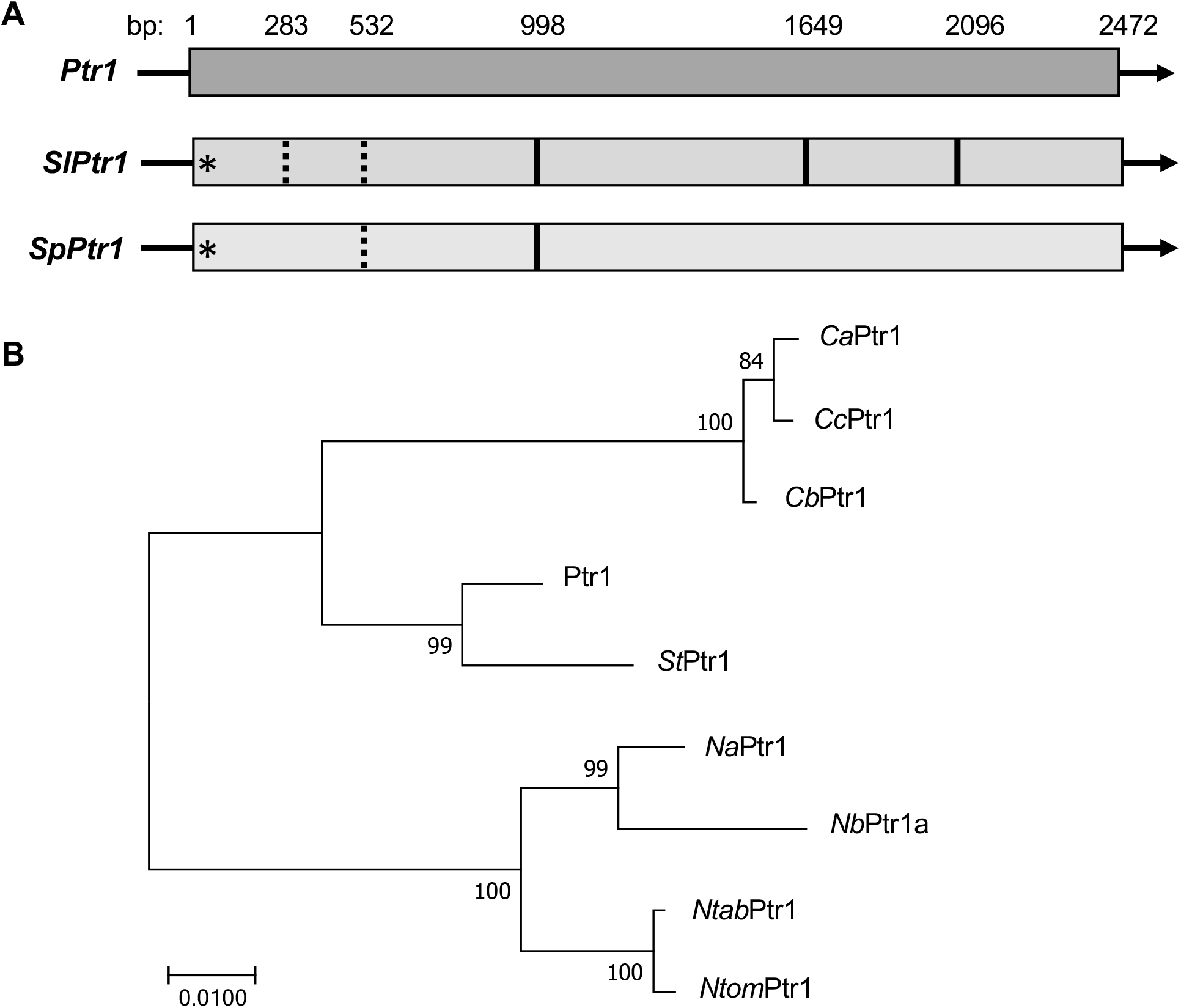
*Ptr1* orthologs in various solanaceous species. **A)** Schematic of *Ptr1* gene structure in *S. lycoperiscoides* (*Ptr1*), *S. lycoperiscum* (*SlPtr1*), and *S. pennellii* (*SpPtr1*). *Ptr1* has a single exon (gray bar) of 2,472 base pairs. *SlPtr1* and *SpPtr1* are pseudogenes that lack a start codon (asterisk) and have other nonsense mutations (dotted bars) and frameshift mutations (solid bars). The position of each mutation is indicated relative to the *Ptr1* base pair (bp) position. **B)** Ptr1 is conserved in many species in the Solanaceae family. Maximum likelihood tree generated from the amino acid sequences of *S. lycopersicoides* Ptr1 and homologs from *S. tuberosum (St)*, *Capsicum chinese (Cc), Capsicum bacctum (Cb), Capsicum annum (Ca)*, *Nicotiana tomentosiformis (Ntom), Nicotiana tabacum (Ntab), Nicotiana benthamiana (Nb)*, and *Nicotiana attenuate (Na)*. Bootstrapping 1000 replicates, with bootstrap values over 75 shown. The tree is drawn to scale, with branch lengths measured in the number of substitutions per site.

A broader search identified intact *Ptr1* orthologs in a variety of other solanaceous plants, including potato, three species of pepper, and four species of tobacco including *N. benthamiana* (**Figure 3B**). In all of these species the predicted amino acid similarity between Ptr1 and that encoded by the orthologs is greater than 95% (**Figure S5** and **Table S3**).

### The *Ptr1* ortholog in *N. benthamiana* and in potato mediates recognition of AvrRpt2 and RipBN

The observation that expression of AvrRpt2 alone in *N. benthamiana* leaves causes cell death raised the possibility that the *Ptr1* ortholog in this species is responsible for this response. As expected for an allotetraploid, *N. benthamiana* has two *Ptr1* orthologs (*NbPtr1a* and *NbPtr1b*), although a 5-base pair deletion in *NbPtr1b* leads to a premature stop codon (**Figure S6**). To test whether *NbPtr1a* plays a role in AvrRpt2-mediated cell death in *N. benthamiana*, we selected a 325-bp fragment of this gene for use in tobacco rattle virus (TRV)-induced gene silencing (VIGS; **Figure S6**). *N. benthamiana* (Nb1; (Bombarely *et al*., 2012)) plants were infected with TRV:*Ptr1* or a control, TRV:*EC1*, containing a small fragment of *E. coli* DNA (Rosli *et al*., 2013). Expression of AvrRpt2 or RipBN, but not of the controls AvrRpt2(C122A) or YFP, induced strong cell death in uninfected Nb1 leaves, as well as those infected with TRV:*EC1* (**Figure 4A**). No cell death was observed in *NbPtr1*-silenced plants, indicating the *Ptr1* ortholog mediates AvrRpt2 recognition in *N. benthamiana*. The constitutive-active Prf protein induced cell death in all the plants as expected.

**Figure 4.**
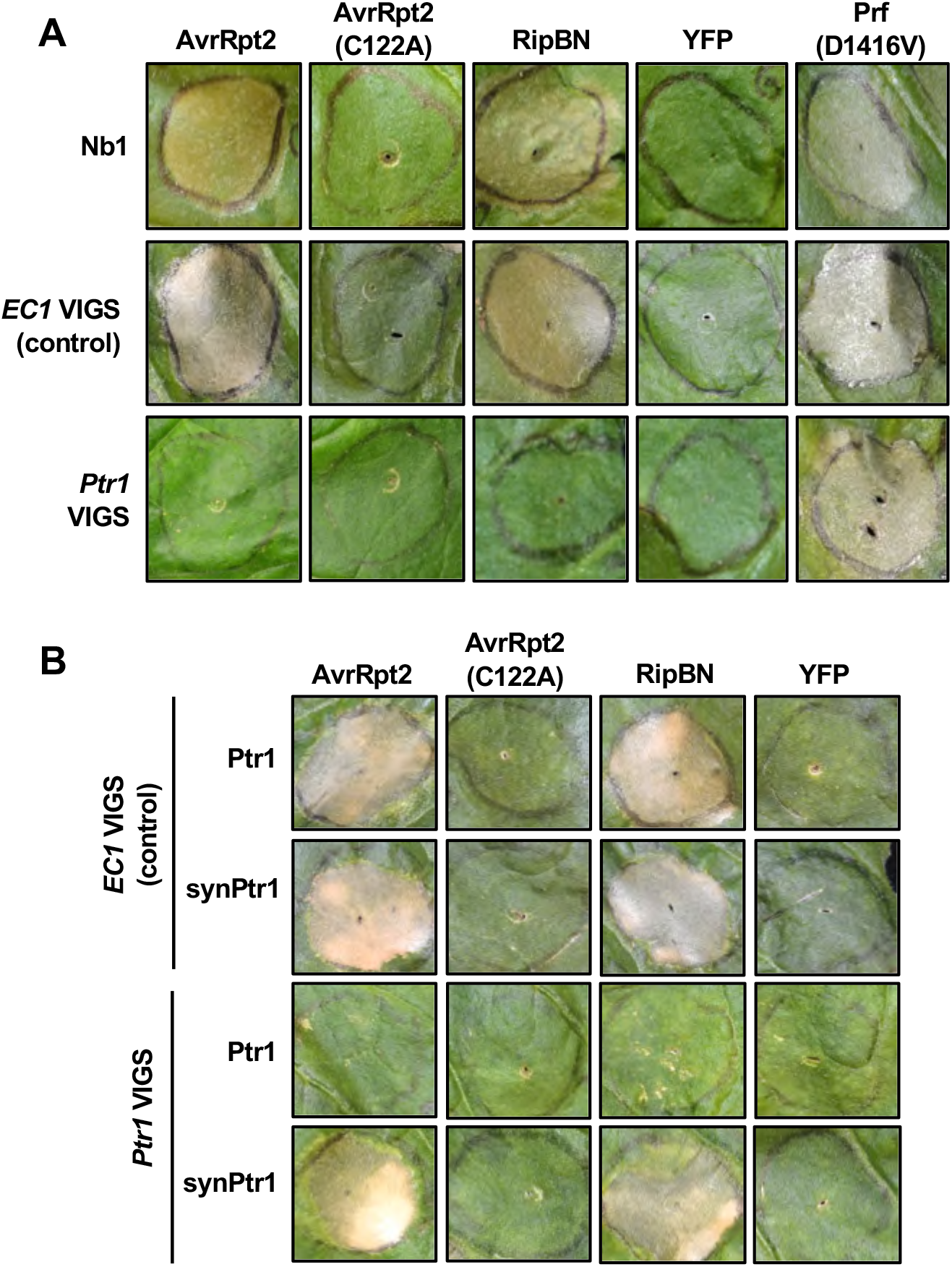
The *Ptr1* homolog in *N. benthamiana* mediates cell death response to AvrRpt2 and RipBN. **A)** *Agrobacterium* strains carrying *AvrRpt2:Myc* (OD_600_ = 0.2), *AvrRpt2(C122A):Myc* (OD_600_ = 0.2), *RipBN:Myc* (OD_600_ = 0.2), *YFP:Myc* (OD_600_ = 0.2), or *35S:Prf(D1416V):HA* (OD_600_ = 0.2) were infiltrated into leaves of *N. benthamiana* (Nb1) or Nb1 plants silenced with either TRV:*Ptr1* (*Ptr1* VIGS) or TRV:*EC1* (*EC1* VIGS). Photographs were taken 48 hr after agroinfiltration (except *35S:Prf(D1416V):HA)* in Nb1 leaves and after 72 hr for VIGS leaves. *35S:Prf(D1416V):HA* treatment was photographed 4 days after infiltration for both Nb1 and VIGS plants. See Figure S6 for details of *NbPtr1* VIGS construct. Photographs are representative of three independent experiments. **B)** A *Ptr1* gene was synthesized (*synPtr1*) with a divergent nucleotide sequence to avoid silencing but retaining the same amino acid sequence (Figure S7). *Agrobacterium* strains harboring AvrRpt2, AvrRpt2(C122A), RipBN, and YFP were infiltrated into leaves at OD_600_ = 0.05 co-expressed with either *35S:Ptr1:HA* or *35S:synPtr1:HA* (both at OD_600_ = 0.025). Photographs were taken 6 days after infiltration and are representative of three independent experiments. All genes were expressed from the CaMV 35S promoter.

To assure that the loss of AvrRpt2-induced cell death we observed in the *Ptr1*-silenced plants was not due to silencing of an ‘off-target’ gene, we developed a synthetic version of *Ptr1* (*synPtr1*) with a divergent DNA sequence that would make it resistant to silencing, yet encode an identical amino acid sequence (**Figure S7**). Immunoblotting showed that both Ptr1 and synPtr1 proteins accumulated in the TRV:*EC1* control plants, whereas in *Ptr1*-silenced plants only the protein encoded by *synPtr1* accumulated (**Figure S8**). Co-expression in *Ptr1*-silenced plants of synPtr1 with either AvrRpt2 or RipBN induced cell death whereas cell death did not occur in silenced plants co-expressing Ptr1 with the effectors (**Figure 4B**). These experiments demonstrate that Ptr1 and its function is conserved in *N. benthamiana*.

The protein predicted to be encoded by the *Ptr1* ortholog in potato (*StPtr1*) is 97% identical to the Ptr1 protein and it seemed likely that it might also mediate recognition of AvrRpt2 and RipBN (**Figure S5**). Attempted use of *Agrobacterium*-infiltration to transiently express AvrRpt2 or RipBN in potato leaves did not cause cell death possibly due to the inefficiency of this method in this species. We therefore cloned the *StPtr1* ortholog from potato variety Dakota Crisp and expressed it using a high titer of *Agrobacterium* in *N. glutinosa* leaves. *St*Ptr1 caused cell death on its own at this titer which was inhibited by co-expression of *Sl*Rin4-3 just as observed for Ptr1 (**Figure 5A**). As with Ptr1, using a lower titer of *Agrobacterium* prevented *St*Ptr1-induced cell death and co-expression of AvrRpt2 or RipBN in this condition induced cell death (**Figure 5B**). *Ptr1* is therefore conserved in diverse solanaceous species, but is a pseudogene in tomato and *S. pennellii*.

**Figure 5.**
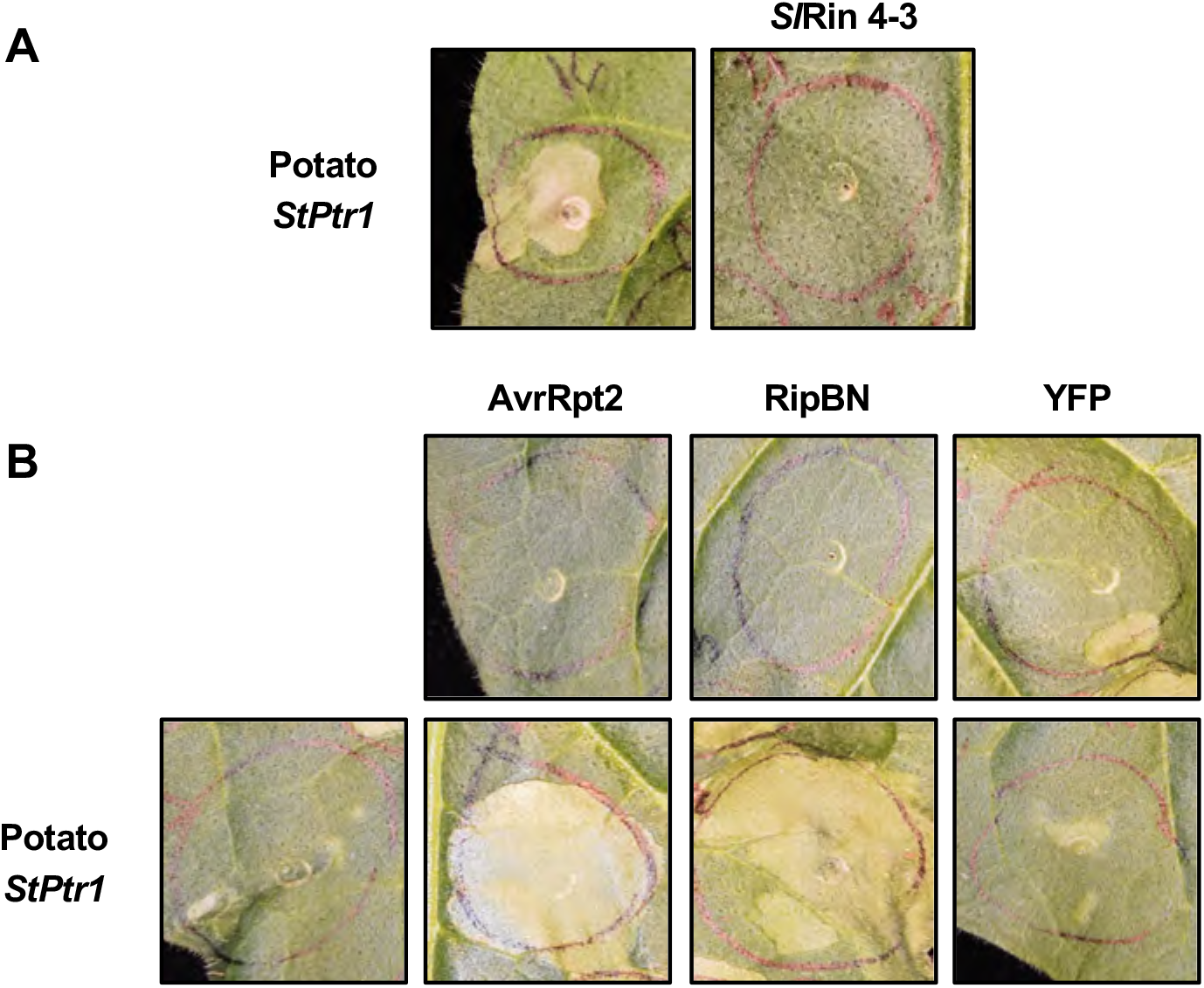
The *Ptr1* ortholog in potato (*StPtr1*) mediates recognition of AvrRpt2 and RipBN. **A)** *Nicotiana glutinosa* leaves were syringe-infiltrated with *Agrobacterium* strains carrying the potato *Ptr1* ortholog (OD_600_ = 0.1) and *Myc:SlRin4-3* (OD_600_ = 0.2) both expressed from the CaMV 35S promoter. Photographs were taken 48 hr after agroinfiltration. **B)** *N. glutinosa* leaves were syringe-infiltrated with *Agrobacterium* strains carrying the potato *Ptr1* ortholog (OD_600_ = 0.025), *AvrRpt2:Myc* (OD_600_ = 0.05), *RipBN:Myc* (OD_600_ = 0.025), and *YFP:Myc* (OD_600_ = 0.05) with all genes expressed from the CaMV 35S promoter. Photographs were taken 41 hr after agroinfiltration. Data shown are representative of three independent experiments, using three biological replicates and infiltrating two leaves per plant.

### *Ptr1* is not orthologous to *Mr5* or *RPS2*

The nucleotide sequence of the *Ptr1* gene is 39% identical to *RPS2* and 34% identical to *Mr5*, which mediate recognition of AvrRpt2 in Arabidopsis and apple, respectively (**Figure S9**). The amino acid sequence identity is similarly low (~20% identity of the Ptr1 protein with either RPS2 or Mr5) with many of the identical residues being in conserved NLR motifs (**Figure S10**). To understand the broader relationships of the *Ptr1*, *RPS2*, and *Mr5* genes we searched the *S. lycopersicoides* genome sequence to find the most closely related genes for each of them. The NB-ARC domains encoded by these genes were then used in a phylogenetic analysis. This revealed that multiple *S. lycopersicoides* proteins are more similar to RPS2 and Mr5 than to Ptr1 with each of the proteins belonging to a distinct clade (**Figure 6**). A similar analysis looking for the most closely related genes in Arabidopsis and in apple confirmed that multiple genes occur in these two species which are closer to *Ptr1* than to either *RPS2* or *Mr5* (**Figure S11**).

**Figure 6.**
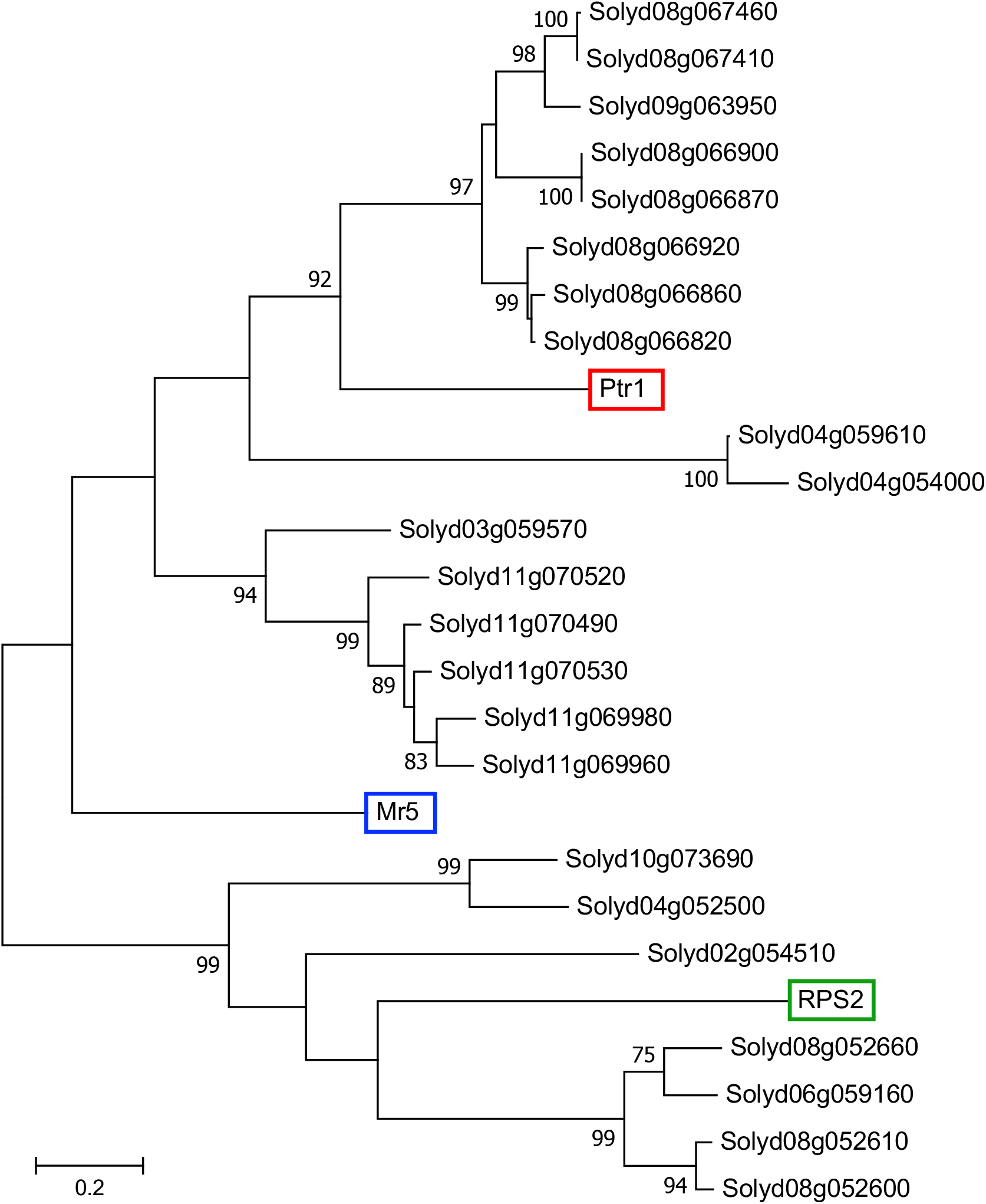
Phylogenetic tree of Ptr1, Mr5, RPS2 with their most closely related proteins in *S. lycopersicoides*. Maximum likelihood tree of the amino acid sequence of the NB-ARC domain only of Ptr1, Mr5, and RPS2 top BLAST hits with an intact NB-ARC domain, including the P-loop, kinase 2, kinase 3a, and GLPL motifs, from the *S. lycopersicoides* genome protein database. 1000 bootstrap replicates, with bootstraps 75 or above shown. The tree is drawn to scale, with branch lengths measured in the number of substitutions per site.

## Discussion

A natural outbreak of bacterial speck disease led to the serendipitous discovery of the *Ptr1* locus which confers resistance to *Pst* strains that express the effector AvrRpt2 (Mazo-Molina *et al*., 2019). The current annotation of the *S. lycopersicoides* genome sequence indicated there are 16 NLR-encoding genes in the large introgression segment shared between LA4245-R and LA4277-R and we hypothesized that one of them is *Ptr1*. Eight of these NLR-encoding genes are clustered within a ~2 Mb region while the other eight are dispersed in an almost 80 Mb segment of chromosome 4. As expected when working with a wild relative of tomato, we observed little overall recombination involving the introgressed region. However, a single recombinant among the 585 F2 plants screened eliminated the eight clustered NLR-encoding genes as being candidates for *Ptr1*. Inspection of the gene models for the remaining eight genes revealed one of them to be a pseudogene and no transcripts were detectable in leaves of five of the remaining genes, leaving just two candidates. *Agrobacterium*-mediated transient expression assays revealed that one of these two genes is *Ptr1* and showed that it mediated recognition of both AvrRpt2 and RipBN via their ability to cause degradation of Rin4. Here we discuss the use of *N. glutinosa* in this work, the discovery of *Ptr1* homologs in various solanaceous species, potential use of *Ptr1* in breeding programs and the process of convergent evolution which apparently is responsible for the ability of three diverse NLR-encoding genes to mediate recognition of AvrRpt2.

It was reported previously that AvrRpt2 does not induce cell death in *N. glutinosa* (Kessens et al., 2014). We therefore used this species for *Agrobacterium*-mediated transient expression of the *Ptr1* candidates because in our conditions AvrRpt2 induces strong cell death in *N. benthamiana*, the typical species used for agroinfiltration. No genome sequence is currently available for *N. glutinosa* but it appears to lack a functional *Ptr1* gene since our experiments show that the Ptr1 pathway is otherwise intact in this species. *N. glutinosa* does appear to have one or more functional Rin4 proteins because at a lower titer Ptr1-induced cell death does not occur, whereas co-expression of AvrRpt2 at this low titer (likely leading to degradation of Rin4) induces cell death. Higher expression of Ptr1 by using higher titers of *Agrobacterium* causes cell death without AvrRpt2 likely because it disrupts the stoichiometric levels of Ptr1 and Rin4 needed to negatively regulate Ptr1. Indeed, as shown previously with RPS2, which also causes cell death on its own, we found that co-overexpression of Rin4 and Ptr1 suppressed Ptr1 autonomously-induced cell death.

*Arabidopsis* RIN4 has two sites (RCS1 and RCS2) each with a consensus sequence which is proteolytically cleaved by AvrRpt2, resulting in three AvrRpt2-cleavage products: ACP1, ACP2, and ACP3 (Coaker *et al*., 2005). The C-terminal half of *At*RIN4 is necessary and sufficient for the negative regulation of RPS2, but none of the individual cleavage products alone can negatively regulate RPS2. This supports a model that cleavage of RIN4 specifically at RCS2 leads directly to RPS2 activation via loss of suppression by RIN4 (Day *et al*., 2005, Kim *et al*., 2005). Interestingly, different from *At*RIN4, apple *Md*RIN4 does not suppress NLR-dependent autoactivity. Instead, ACP3 released upon cleavage of *Md*RIN4 is sufficient to activate Mr5 (Prokchorchik *et al*., 2019). Investigation of the mechanism by which ACP3 from *Md*RIN4, and not *At*RIN4, activates Mr5 revealed two polymorphic amino acid residues in the N-terminal sequences of ACP3 in *At*RIN4 (N158/Y165) and *Md*RIN4 (D186/F193) (Prokchorchik *et al*., 2019). The three tomato Rin4 proteins that are expressed in leaves each have a hybrid of these polymorphisms. That is, instead of having an asparagine (N) as occurs in *Arabidopsis*, *Sl*Rin4-1, 4-2, and 4-3 have an aspartic acid (D) like apple. Moreover, the tomato Rin4 proteins have a tyrosine (Y) like *At*RIN4 instead of a phenylalanine (F) as occurs in *Md*RIN4 (**Figure S1A**). Whether or not these differences play a role in activation or suppression of Ptr1 is unknown and their future investigation may shed light on how the SlRin4 proteins regulate Ptr1.

Despite much effort, by many researchers, no source of simply-inherited resistance to race 1 strains of *Pst* has been discovered among accessions of cultivated tomato or its wild relatives (although some QTLs contributing to race 1 resistance have been reported; (Bao *et al*., 2015, Thapa *et al*., 2015, Hassan *et al*., 2017)). The discovery that the *Ptr1* ortholog in both tomato and *S. pennellii* has multiple mutations including one that disrupts the start codon explains why race 1 resistance involving recognition of AvrRpt2 was never found. This observation suggests that strains of *Pst* expressing AvrRpt2 were not serious pathogens in the environment where wild relatives of tomato evolved and that there was no selection pressure to retain *Ptr1*. It will be interesting in the future to determine if all or just some of the accessions of the 12 wild relatives of tomato have nonfunctional versions of *Ptr1* and possibly correlate this feature with the regions in which the accessions were originally collected.

The *Ptr1* ortholog from potato was also able to mediate recognition of AvrRpt2 and RipBN when transiently expressed in *N. glutinosa* leaves. We cloned this ortholog from the variety Dakota Crisp and our version is 98% similar to a gene annotated in the potato genome sequence as resistance gene analog 1 (*RGA1*; GenBank No. XP_006340095.1). No function for the *RGA1* gene has been reported and we have therefore named it *StPtr1*. The presence of a functional *Ptr1* gene in both potato and *S. lycopersicoides*, both of which originated in South America, and *N. benthamiana* which originated in Australia, supports the hypothesis that bacterial pathogens of these diverse solanaceous species and their progenitors have, over evolutionary time, expressed an AvrRpt2-like protein. This is consistent with the importance of this effector for bacterial virulence and with its presence in many bacterial species and multiple pathovars of *P. syringae* (**Figure S12**).

The observation that expression of AvrRpt2 or RipBN alone causes cell death in *N. benthamiana* leaves and the discovery of an intact *Ptr1* ortholog (*NbPtr1a*) in this tobacco species raised the possibility that the effector-induced cell death involves *Ptr1*. Virus-induced gene silencing of the *NbPtr1a* gene in *N. benthamiana* and complementation with a synthetic *Ptr1* gene that is recalcitrant to silencing confirmed this is the case. *N. benthamiana* has been used extensively to characterize biochemical aspects of the Pto/Prf complex and to identify and study host proteins that play a role in the Pto/Prf pathway such as Cbl10, Cpk6, Epk1, Hsp90, Mai1, Nrc2/3, MAPKKK*α*, MKK2, Sgt1, TFT7 (Ekengren *et al*., 2003, Lu *et al*., 2003, del Pozo *et al*., 2004, Mucyn *et al*., 2009, Oh *et al*., 2010, Oh and Martin, 2011, de la Torre *et al*., 2013, Kud *et al*., 2013, Ntoukakis *et al*., 2013, Saur *et al*., 2015, Wu *et al*., 2017, Roberts *et al*., 2019, Wu and Kamoun, 2019). It will be interesting in the future to determine if all of these proteins play a role in the Ptr1 pathway or if the Pto/Prf and Ptr1 pathways use some divergent host components.

We have considered previously how the *Ptr1* locus could play an important role in protection of tomato against bacterial speck disease (Mazo-Molina *et al*., 2019). The identification of the *Ptr1* gene itself will now allow it to be efficiently tracked as it is backcrossed into various breeding lines. *Ptr1* and *Pto* are located on different chromosomes (4 and 5, respectively) which will facilitate the development of tomato varieties containing both genes. Such varieties would confer resistance to all currently known races of *Pst*. The low rate of recombination between *S. lycopersicoides* and tomato DNA could interfere with introgression of a small segment of *S. lycopersicoides* carrying the *Ptr1* gene. Methods are available to address low recombination such as using a bridge species and we have discussed these previously (Mazo-Molina *et al*., 2019). Finally, if the new method of CRISPR ‘Prime editing’ proves to be feasible in tomato then it might be employed to ‘repair’ the *Ptr1* pseudogene present in tomato (Anzalone *et al*., 2019). Such an approach would greatly simplify the introgression of the modified gene into advanced breeding lines for use in development of speck-resistant tomato varieties.

By using a phylogenetic analysis we found that the most similar genes to *RPS2* and *Mr5* in *S. lycopersicoides* fall into distinct clades which are distantly related to *Ptr1*. The same conclusion came from phylogenetic trees of the most closely related genes to *Ptr1* in the apple and Arabidopsis genomes. These observations indicate that *Ptr1*, which is present in at least three solanaceous species, is not orthologous to either *RPS2* or *Mr5* and likely arose by convergent evolution to mediate recognition of AvrRpt2. *Ptr1*, *RPS2* and *Mr5* represent just the third case where non-orthologous NLR-encoding genes in different plant species have been reported to mediate recognition of the same effector. In soybean, the *R* genes *Rpg1b* and *Rpg1r* recognize AvrB and AvrRpm1, respectively, whereas in *Arabidopsis* RPM1 recognizes both of these effectors. Although all of these genes encode CC-NLR proteins, which detect alteration of RIN4, the *Rpg1* genes are not orthologous to *RPM1* (Ashfield *et al*., 2004). The other example is the recent discovery of an NLR-encoding gene in barley, *Pbr1*, whose protein detects AvrPphB-directed cleavage of PBS1 as does the *Arabidopsis* RPS5 protein; the *RPS5* and *Pbr1* genes are not orthologous (Carter *et al*., 2018). These three examples demonstrate the remarkable plasticity inherent in the modular structure of NLR proteins which facilitates the generation of common resistance specificities from highly divergent genes.

The specific mechanisms by which RPS2 and Mr5 are activated upon degradation of Rin4 are unknown (Prokchorchik *et al*., 2019, Toruño *et al*., 2019). The availability now of amino acid sequences of three diverse proteins that mediate recognition of AvrRpt2 might provide insights into subdomains of the proteins that are involved in the response to AvrRpt2 (and to Rin4 cleavage products). As expected, there are a number of residues in common between the Ptr1, *St*Ptr1, *Nb*Ptr1a, RPS2, and Mr5 proteins in the NB-ARC domain (**Figure S10**). More interesting perhaps is the presence of some conserved residues in the LRR domain which is typically more divergent among NLR proteins. Whether these conserved residues play a role in mediating the response to AvrRpt2 will be a focus of future work using the *Agrobacterium* infiltration assay in *N. glutinosa*.

## Experimental Procedures

### Bacterial strains

*Pseudomonas syringae* pv. tomato strains JL1065 (Whalen *et al*., 1991) and JL1065Δ*avrRpt2* (Lim and Kunkel, 2005) were grown on Kinǵs B (KB) semi-selective media at 30°C. *Agrobacterium tumefaciens* strains GV3101 and GV2260 (Holsters *et al*., 1980) were grown on LB with appropriate antibiotics at 30°C (Table S5). All strains were stored in 20% glycerol + 60 mM sucrose at –80°C. *Escherichia coli* was used for plasmid maintenance and grown in LB medium at 37°C.

### Plant materials

Seeds of *Solanum lycopersicoides* introgression lines were obtained from the Tomato Genetics Resource Center (https://tgrc.ucdavis.edu/lycopersicoides_ils.aspx). LA4245-R and LA4277-R are maintained as heterozygotes (*Ptr1/ptr1*). Progeny derived from selfing these plants are rarely homozygous *Ptr1/Ptr1* (around 3% of the progeny) and such plants grow more slowly than *Ptr1/ptr1* or *ptr1/ptr1* plants. Tomato plants were grown in a greenhouse at 24°C during daylight and 22°C at night. *Nicotiana benthamiana* Nb1 (Bombarely *et al*., 2012) and *Nicotiana glutinosa* plants were maintained in a growth chamber with 16 hr:8 hr, light:dark at 24°C with light and 20°C in the dark and 50% humidity. Tomato and *Nicotiana* plants were grown in Cornell Osmocote Mix soil (0.16 m^3^ peat moss, 0.34 m^3^ vermiculite, 2.27 kg lime, 2.27 kg Osmocote Plus15-9-12 and 0.54 kg Uni-Mix 11-5-11; Everris, Israeli Chemicals Ltd). After pathogen inoculation, tomato plants were moved to a growth chamber with 25°C, 50% humidity, and 16 hr light.

### Mapping of *Ptr1*

DNA from progeny of selfed LA4277-R (*Ptr1/ptr1*) plants was isolated with DNA extraction buffer (200 mM Tris-HCl pH 8.0, 250 mM NaCl, 25 mM EDTA pH 8, 0.5% w/v sodium dodecyl sulfate) and resuspended in distilled water. Simple sequence repeat (SSR) markers were designed by mapping the *S. lycopersicoides* genome to *S. lycopersicum* Heinz 1706 v 3.0 using nucmer 4.0.0beta (Delcher *et al*., 1999, Marcais *et al*., 2018). A pipeline was developed to identify indels between 30 - 200 bp and develop flanking primers in order to amplify PCR products of different sizes based on insertions or deletions (Untergasser *et al*., 2012)(**Table S6**). Potential recombinants were selfed and the resulting progeny were genotyped to confirm the segregating recombination event. Recombinant progeny were phenotyped by vacuum infiltration with *Pst* JL1065 or *Pst* JL1065Δ*avrRpt2* as described previously (Mazo-Molina *et al*., 2019) and visually monitored for the absence or presence of disease symptoms.

### RNA-Seq

Seven-week-old LA4277-Ro (*Ptr1/Ptr1*) plants were identified by markers and vacuum-infiltrated with a suspension of *Pst* JL1065 at 2 x 10^7^ cfu/mL. Four biological replicates were performed for each treatment. Tissue samples were collected at 5 hr after infiltration. Total RNA was isolated using RNeasy Plant Mini Kit (Qiagen) with additional in-column DNase digestion using the RNase-Free DNase Kit (Qiagen). Libraries for 3’ RNA-Seq were prepared by the Cornell Biotechnology Resource Center’s Genomics Facility using the Quantseq FWD kit (Lexogen).

Raw RNA-Seq reads were processed, removing adaptors and low-quality sequences using Trimmomatic (Bolger *et al*., 2014b). Low quality sequences were removed from leading and trailing read ends, and trimmed reads shorter than 10 bases were discarded. Clean reads were then aligned to the SILVA rRNA database (Quast *et al*., 2013) using Bowtie (Langmead and Salzberg, 2012) allowing for up to three mismatches. Reads that mapped to rRNA sequence were discarded. The final high-quality reads for each library were aligned to the *S. lycopersicoides* LA2951 reference genome (https://solgenomics.net/organism/Solanum_lycopersicoides/genome) using STAR default parameters (Dobin *et al*., 2013). Raw counts for each LA2951 gene model were generated by counting the total number of reads that mapped between the gene region and 500 bp downstream of its stop codon. Reads were normalized to reads per million (RPM).

### Cloning of *Ptr1* candidates and tomato Rin4 genes

*Ptr1* candidate genes were cloned from LA4277-Ro (*Ptr1/Ptr1*) cDNA into pBTEX (**Table S7**) using the In-fusion cloning kit manufacturer’s instructions (Takara). *StPtr1* was cloned from Dakota Crisp potato plants (*Solanum tuberosum*) into pBTEX via the In-fusion cloning kit (Takara). Nucleotide and amino acid sequences have been deposited in GenBank for *Ptr1* from *S. lycopersicoides* (GenBank accession no. MT134103), *N. benthamiana*, *NbPtr1a* (MT134102) and potato *StPtr1* (MT134101)(**Table S3**).

*Sl*Rin4 genes were cloned from LA4245 cDNA into pJLSmart (Mathieu *et al*., 2014) and recombined into the Gateway expression vectors pGWB518 and pGWB417 (Nakagawa *et al*., 2007) using LR Clonase II according to the manufacturer’s instructions (Thermo Fisher Scientific) to generate N-tagged and C-tagged proteins, respectively.

Constructs were transformed into *E. coli* Stellar competent cells (Clontech). Inserts of all plasmids were sequenced from *E. coli* and then transformed into *Agrobacterium tumefaciens* GV2260. See **Table S6** and **Table S7** for a list of all oligo and constructs used in this study.

### LA4277 genomic DNA library preparation and Oxford Nanopore sequencing

The LA4277 genome sequence data are available at NCBI as BioProject No. PRJNA610286. To extract high molecular weight DNA, nuclei were enriched from 2 g of LA4277-Ro (*Ptr1/Ptr1*) leaves using a method modified from (Gendrel *et al*., 2005). Two g of leaf tissue was harvested and ground in liquid nitrogen and incubated with 90 mL of 0.4 M sucrose, 10 mM Tris-HCl pH 8, 10 mM MgCl_2_, 5 mM BME (β-mercaptoethanol) for 10 min on ice with shaking. Large leaf debris was removed by filtering through 8 layers of filter paper (Fisher Scientific) and one layer of miracloth (Fisher Scientific). The solution was centrifuged at 3000*xg* at 4°C for 20 min. The resulting pellet was resuspended in 500 uL of 0.25 M sucrose, 10 mM Tris-HCl pH 8, 10 mM MgCl_2_, 1% Triton X-100, 5 mM BME. The suspension was then centrifuged at 12,000*xg* at 4°C for 10 min. The pellet was resuspended in 300 uL 0.25 sucrose, 20 mM Tris-HCl pH 8, 4 mM MgCl_2_, 0.3% Triton X-100 and 500 uL of 2.5 M sucrose was then mixed with the nuclei suspension. This mixture was then overlayed on top of 800 uL of 1.7 M sucrose, 10 mM Tris-HCl pH 8, 2 mM MgCl_2_, 0.15% Triton X-100, 5 mM BME and centrifuged at 16,000*xg* at 4°C for 1 hr. The pelleted nuclei were washed 2x in 500 uL of 25% glycerol, 20 mM Tris-HCl pH 7.4, 2.5 mM MgCl_2_, 0.2% Triton X-100. DNA was isolated from the enriched nuclei suspension as described previously (Bernatzky and Tanksley, 1986).

Covaris g-tube was used in accordance with the manufacturer’s protocol to shear 8 ug DNA to 20 kb fragments. Following shearing, small fragments were removed with a size-selection step using 1x AMPure XP (Beckman Coulter, Brea, CA, USA) in NaCl/PEG buffer (10 mM Tris-HCl, 1 mM EDTA pH 8, 1.6 M NaCl, 0.25% Tween-20, 11% PEG-800) (Nagar and Schwessinger, 2018). Large fragments were eluted from beads in 50 uL H_2_O. A library was prepared using 1 ug of DNA as input using the SQK-LSK109 kit (Oxford Nanopore Technologies, Oxford, UK) according to manufacturer’s protocol, except incubation times were increased to 30 min for DNA repair, end-prep and 1 hr for adapter ligation, and 10 min for all elutions from beads. The finished library was loaded on to a single FLO-Min106D R9 Spot-ON flowcell (Oxford Nanopore Technologies, Oxford, UK). MinION sequencing was performed using the standard script for a 60 hr run.

### LA4277 genomic DNA library preparation and Illumina sequencing

Illumina libraries for whole genome sequencing of LA4277-Ro were prepared as described previously (Mazo-Molina *et al*., 2019) using nuclear-enriched DNA. Paired-end 150-bp DNA reads were sequenced using the Illumina HiSeq 2000 platform by Genewiz (South Plainfield, NJ)

### LA4277 genome assembly

Nanopore sequence was base-called using Guppy v 2.3.5+53a111f (Wick *et al*., 2019). A hybrid approach implemented in MaSuRCA v3.3.2 (Zimin *et al*., 2013) was used to de novo assemble the LA4277 Nanopore and Illumina genome sequence. The assembly was polished with Illumina reads and 3 rounds of Pilon correction. The assembly had a total length of 840.2 Mbp, N50 of 1.3 Mbp, and captured over 97% of the BUSCO set.

#### *Agrobacterium*-mediated transient protein expression

Cell death assays in five-week-old *N. benthamiana* and *N. glutinosa* plants were performed as described previously (Oh *et al*., 2010), after dilution of the *Agrobacterium* strains to a final OD_600_ of 0.025 or 0.1 as indicated in the figure legend for Ptr1, 0.05 for AvrRpt2, 0.2 for Rin4, and 0.1 for Prf. Cell death started to appear 2 days after *Agrobacterium* infiltration, except for Prf where it started 4 days after infiltration. To detect protein expression in *N. benthamiana*, a final OD_600_ of 0.1 for Ptr1, 0.05 for AvrRpt2, and 0.2 for Rin4 was used and leaf tissue was sampled 28 hr after infiltration.

For the VIGS experiments, *Agrobacterium* strains were diluted to a final OD_600_ of 0.2 for AvrRpt2, AvrRpt2(C122A), RipBN, YFP, and Prf(D1416V). Cell death started to appear 2 days after infiltration at which time the Nb1 plants were scored and photographed. TRV:*EC1* and TRV:*Ptr1* silenced plants were scored for cell death and photographed 72 hr after infiltration for all constructs except Prf(D1416V), which was scored and photographed 4 days after infiltration. The experiment was repeated three times, using six TRV:*EC1* and TRV:*Ptr1* plants in each experiment and infiltrating two leaves per plant.

For the *synPtr1* complementation assays, *Agrobacterium* containing the Ptr1 and synPtr1 constructs were infiltrated into TRV:*EC1* and TRV:*Ptr1* silenced plants at a final OD_600_ of 0.025 and co-infiltrated with *Agrobacterium* strains carrying AvrRpt2, AvrRpt2(C122A), RipBN, or YFP at a final OD_600_ of 0.05. Cell death started to appear 48 hr after infiltration. Plants were scored for cell death and photographed 6 days after infiltration. The complementation experiment was repeated three times, using 3 - 6 TRV:*EC1* and TRV:*Ptr1* plants in each experiment and infiltrating two leaves per plant. Photos shown in the figures are representative of all replicates. To detect protein expression of Ptr1 and synPtr1 in silenced plants, Agrobacterium strains were infiltrated at a final OD_600_ of 0.1. Leaf samples for protein expression were collected 46 hr after infiltration.

### Immunoblot detection of plant-expressed proteins

To detect protein expression in *N. benthamiana*, total proteins were extracted as previously described (Mazo-Molina *et al*., 2019). To detect AvrRpt2 and Rin4, membranes were probed with anti-c-Myc (GeneScript) antibody at a concentration of 1/7000. Secondary ∝-mouse-HRP was used at a dilution of 1/10,000 (Sigma-Aldrich). For detection of Ptr1, synPtr1, and RPS2 proteins, membranes were probed with anti-HA antibody at a concentration of 1/2,000 (Roche). Secondary ∝-rat-HRP was used at a dilution of 1/10,000 (Cell Signaling Technology).

### Phylogenetic analyses

Alignments were generated using MUSCLE except where noted otherwise. Phylogenetic trees were constructed in MEGA7 (Kumar *et al*., 2016) using the maximum likelihood with a JTT matrix-based model method (Jones *et al*., 1992). Positions containing gaps and missing data were eliminated. A bootstrap analysis with 1,000 replicates was used to determine the confidence probability of each branch (Felsenstein, 1985). Data on which AvrRpt2-like proteins are recognized by Ptr1 are from Mazo-Molina et al. (2019) and the current study.

The top 10 BLASTp hits of the Ptr1, Mr5, and RPS2 NB-ARC domains were identified in the *S. lycopersicoides* v.0.6 genome sequence (Powell *et al*., 2020), *M. domestica* (GDR GDDH13 V1.1) (Jung *et al*., 2019), and *A. thaliana* (TAIR Araport 11) genome protein databases. The NB-ARC domain of each protein hit was determined using Interpro scan (https://www.ebi.ac.uk/interpro/) and the amino acid sequences of the NB-ARC domain for each species were aligned to check for the presence of the NB-ARC conserved motifs. To ensure alignment of the sequences, NB-ARC domains missing any of the motifs were removed from further analysis. AvrRpt2 homologs in Genbank were identified through NCBI BLAST, and were aligned with MUSCLE, along with sequences of *avrRpt2* homologs from recent papers (Eschen-Lippold *et al*., 2016, Dillon *et al*., 2019, Mazo-Molina *et al*., 2019).

### Virus-induced gene silencing (VIGS)

The *Ptr1*-targeting VIGS sequence was selected using the SGN VIGS Tool (Fernandez-Pozo *et al*., 2015). The fragment was cloned into pDONR/Zeo (Invitrogen) followed by an LR reaction (Invitrogen) into pQ11 (Liu *et al*., 2002). The resulting pQ11:*Ptr1* (TRV:*Ptr1*) construct was transformed into *A. tumefaciens* GV2260. The control, pQ11:*EC1* (TRV:*EC1*), contains a small DNA fragment from *E. coli* and was described previously (Rosli *et al*., 2013). VIGS constructs were prepared for infections in *N. benthamiana* as described previously (Chakravarthy *et al*., 2010). Cell death assays were performed five-to-six weeks after agroinfiltration with the VIGS constructs.

### Generation of synthetic *Ptr1*

The synthetic *Ptr1* (*synPtr1*) sequence was designed using the Integrated DNA Technologies (IDT) Codon Optimization Tool (https://www.idtdna.com/CodonOpt) as previously described (Roberts *et al*., 2019) (**Figure S7**). The stop codon was removed in order to generate a C-terminal HA-tag and *Kpn*I and *Stu*I restriction sites were added to the 5’ and 3’ end, respectively, for cloning into pBTEX. The *synPtr1* was synthesized by IDT (Skokie, IL, USA) as a gBlocks fragment and cloned into pBTEX using the In-fusion cloning kit following the manufacturer’s instructions (Takara).

## Author Contributions

Conceptualization: SM, CMM, GBM; Investigation: SM, CMM, BJH, RB, JZ, AF, KS, SRS; Writing (original draft): CMM, SM, GBM; Writing (review and editing): all authors; Funding acquisition: KS, CMM, GBM.

## Acknowledgments

We thank Robyn Roberts and Ning Zhang for helpful comments on the manuscript, Ben Jablonski, Mishi Vachev, Annie Geiger, Christine Kraus, and Sarah Hind for performing supporting experimentation, Zhangjun Fei and Fabio Rinaldi for advice on data analysis and methodology, respectively, Brian Bell and Jay Miller for greenhouse assistance, and Steve McKay for field assistance. Funding was provided by Colciencias Departamento Administrativo de Ciencia, Tecnología e Innovación grant no. 673 (CMM), National Natural Science Foundation of China grant no. 31822046 (KS), and National Science Foundation grant IOS-1546625 (GBM).

---

**Table S1.** Predicted gene models for the *Ptr1* candidates in the *S. lycopersicoides* genome sequence. (https://solgenomics.net/organism/Solanum_lycopersicoides/genome). The start and stop of each gene model corresponds to the genomic coordinates along the chromosome. The number of nucleotides and amino acids correspond to the gene models. Candidate D has a nonsense mutation at base pair 391 which terminates its open reading frame.

| Candidate | Gene ID | Start | Stop | No. Nucleotides | No. Amino Acids |
| --- | --- | --- | --- | --- | --- |
| **A** | Solyd04g059470 | 12054434 | 12051962 | 2472 | 823 |
| **B** | Solyd04g059610 | 12505909 | 12503356 | 2553 | 850 |
| **C** | Solyd04g060430 | 22745706 | 22743765 | 1275 | 424 |
| **D** | Solyd04g060640 | 25255273 | 25252810 | 2464 | pseudogene |
| **E** | Solyd04g061490 | 37884585 | 37886367 | 1782 | 593 |
| **F** | Solyd04g064510 | 74057098 | 74064935 | 2457 | 818 |
| **G** | Solyd04g064750 | 74838450 | 74834836 | 3276 | 1091 |
| **H** | Solyd04g067320 | 80119015 | 80115223 | 3480 | 1159 |

**Table S2.** Transcripts of three of the *Ptr1* candidates are detectable in leaves. 3’ RNA-Seq data (normalized reads per million, RPMs) of the eight *Ptr1* candidates in LA4277-Ro (*Ptr1/Ptr1*) plants that were vacuum infiltrated with *Pst* JL1065 and sampled 5 hr later.

| Candidate | Gene ID | Rep 1 | Rep 2 | Rep 3 | Rep 4 | Mean | Standard deviation |
| --- | --- | --- | --- | --- | --- | --- | --- |
| **A** | Solyd04g059470 | 5.89 | 4.87 | 3.61 | 2.74 | 4.28 | 1.386 |
| **B** | Solyd04g059610 | 0.34 | 1.26 | 0.87 | 0 | 0.62 | 0.558 |
| **C** | Solyd04g060430 | 0 | 0 | 0 | 0 | 0 | 0 |
| **D** | Solyd04g060640 | 1.01 | 0.94 | 1.88 | 1.31 | 1.29 | 0.428 |
| **E** | Solyd04g061490 | 0 | 0 | 0 | 0 | 0 | 0 |
| **F** | Solyd04g064510 | 0 | 0 | 0 | 0 | 0 | 0 |
| **G** | Solyd04g064750 | 0 | 0 | 0 | 0 | 0 | 0 |
| **H** | Solyd04g067320 | 0 | 0 | 0 | 0 | 0 | 0 |

**Table S3.** Details of each *Ptr1* gene in various solanaceous species.

| Species | Gene ID | GenBank ID/NCBI reference sequence | Comments |
| --- | --- | --- | --- |
| *S. lycopersicoides* | Solyd04g059470 | MT134103 | *Ptr1* |
| *S. tuberosum* |  | MT134101 | *StrPtr1*; cv. Dakota Crisp |
| *S. lycopersicum* | Solyc04g039980/Solyc04g039990 |  | no open reading frame |
| *S. pennellii* | Sopen04g016200 |  | no open reading frame |
| *C. annuum* | CA05g00030 | PHT80575.1 | cv. CM334; annotated previously as *RGA1* |
| *N. tabacum* | Nitab4.5_0002085g0050.1 | XP_016435110.1 | gene misannotated. Intron should be exon; correct in GenBank; annotated previously as *RGA3* |
| *N. attenuata* | NIATv7_g26201.t1 | OIT07149.1 | annotated previously as *RGA3* |
| *N. tomentosiformis* | mRNA_17526 | XP_018629570.1 | annotated previously as *RGA3* |
| *C. chinese* |  | PHU16668.1 | hypothetical protein |
| *C. bacctum* |  | PHT54558.1 | hypothetical protein |
| *N. benthamiana* | Niben101Scf07061g00004.1 | MT134102 | *NbPtr1a*; gene misannotated in Nb1 genome sequence. Intron should be exon |
| *N. benthamiana* | Niben101Scf00597g00011.1/Niben101Scf00597g00010.1 |  | *NbPtr1b*; no open reading frame |
| *S. melongena* | SMEL011g377860 |  | missing NB-ARC domain; annotated previously as RPP13-like |
| *P. axillaris* | Peaxi162Scf00970g00162.1 |  | no open reading frame |
| *P. inflata* | Peinf101Scf01668g00022.1 |  | no open reading frame |
| *P. inflata* | Peinf101Scf14222g00007.1 |  | no open reading frame |

**Table S4.**
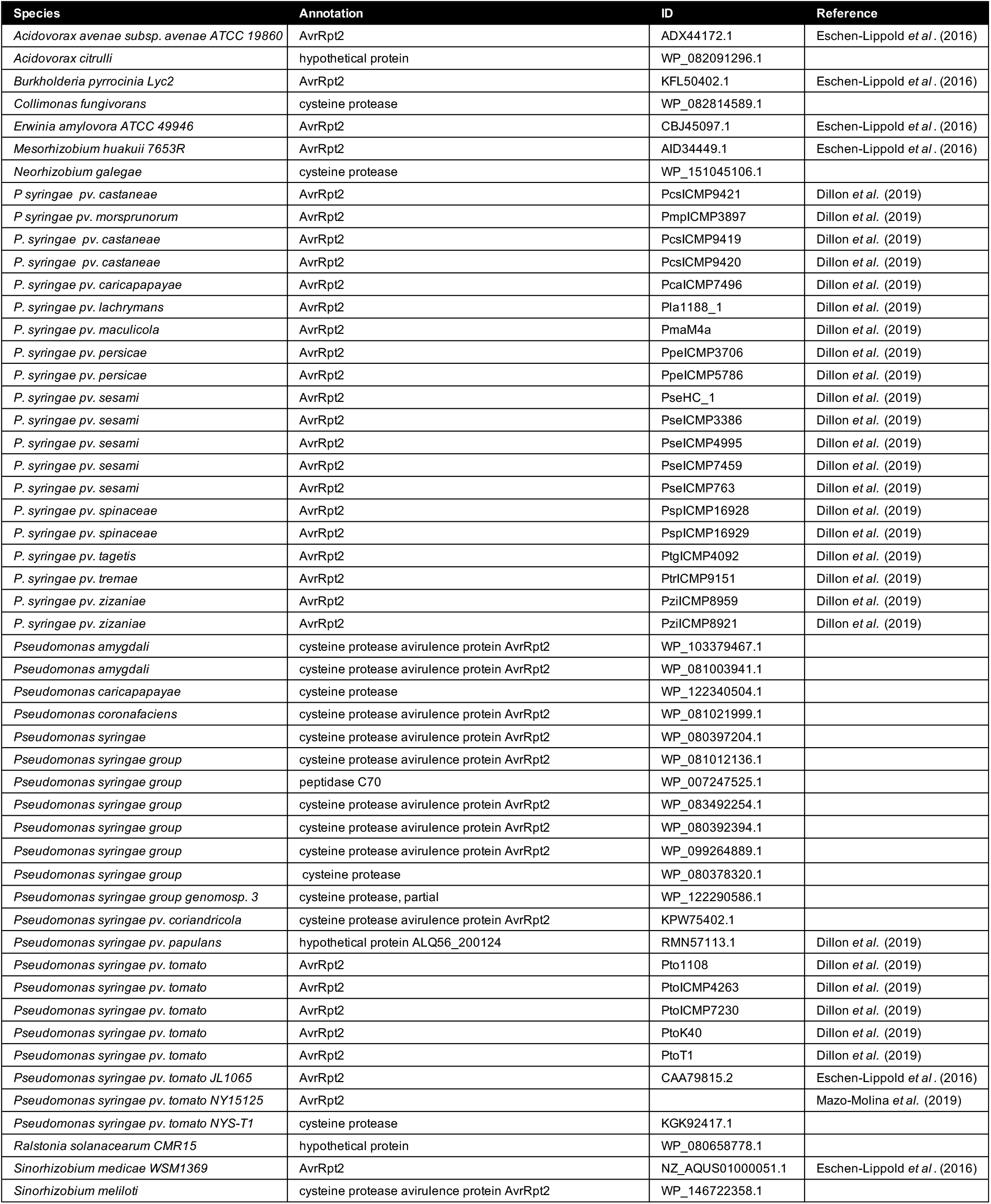
AvrRpt2 sequences used in the phylogenetic tree in Figure S12.

**Table S5:** Bacterial strains used in this study.

| Strain | Genotype | Source | Selection |
| --- | --- | --- | --- |
| *P. syringae* pv. tomato JL1065 | Wild type | Whalen et al. (1991) | Rifampicin |
| *P. syringae* pv. tomato JL1065 | $\Delta avrRpt2$ | Lim and Kunkel et al. (2005) | Rifampicin<br>Spectinomycin |
| *A. tumefaciens* GV3101 + pMP90 | Wild type | Holsters et al. (1980) | Rifampicin<br>Gentamycin |
| *A. tumefaciens* GV2260 | Wild type |  | Rifampicin |
| *E. coli* Stellar | Wild type | Clontech | - |

**Table S6:**
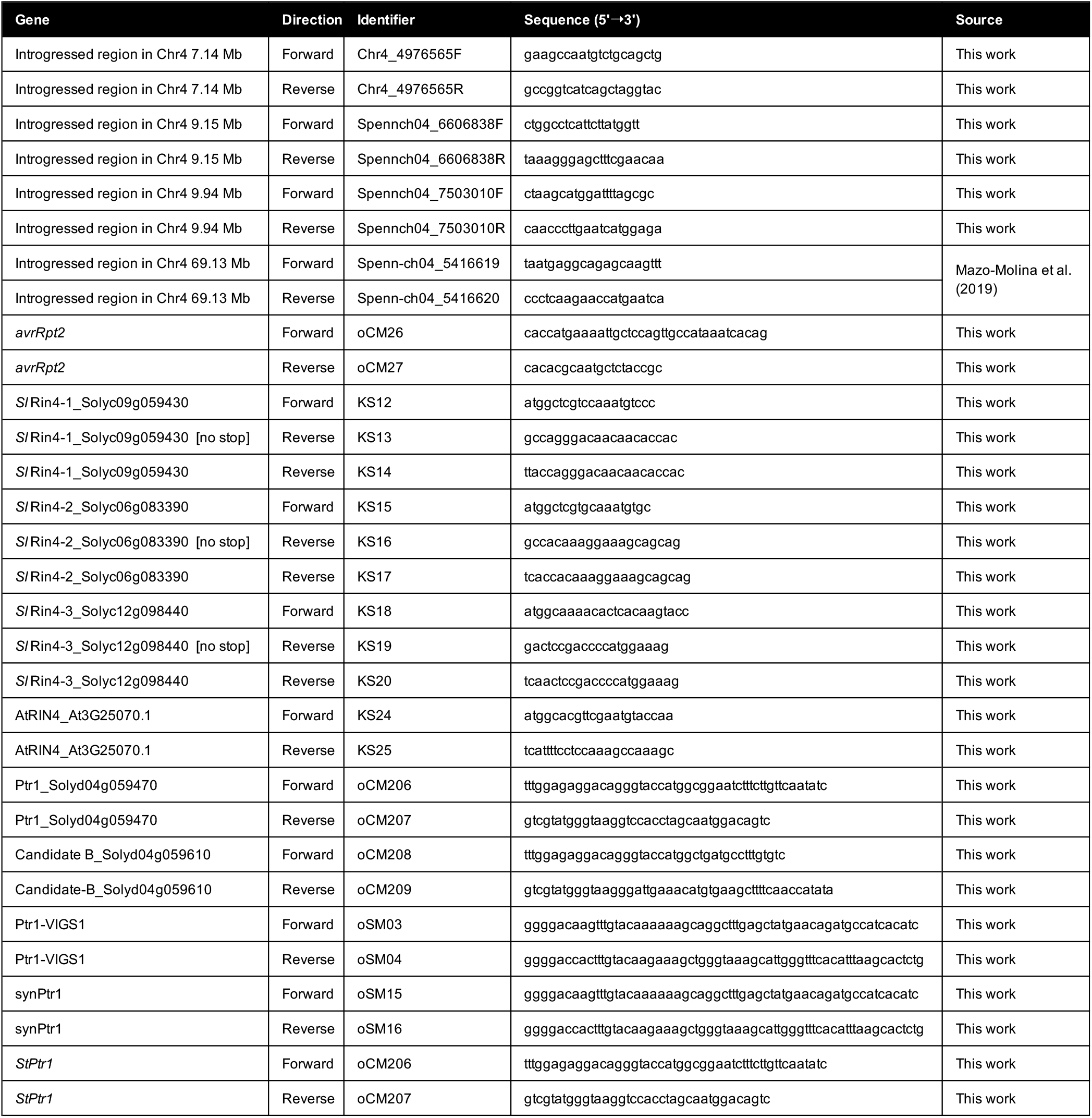
Oligonucleotides used in this study.

**Table S7:** Vectors and plasmids used in this study.

| Vector | Insert | Identifier | Purpose | Source |
| --- | --- | --- | --- | --- |
| pBTEx | - | - | Binary expression vector with N-terminal HA-tag | R. Bressan, Purdue University |
| pBTEx | Ptr1-A_ Solyd04g059470 | pCM64 | Generating N-terminal HA-tag fusion | This work |
| pBTEx | Ptr1-B_ Solyd04g059610 | pCM66 | Generating N-terminal HA-tag fusion | This work |
|  | RPS2_At4g26090 | - | N-terminal HA-tagged RPS2 gene | B. Staskawicz, UC Berkeley |
| pJLSmart | - | - | Binary Gateway entry vector | Mathieu *et al.* (2014) |
| pJLSmart | S/Rin4-1_ Solyc09g059430 | 10347 | Entry clone for N-terminal Myc-tag fusion | This work |
| pJLSmart | S/Rin4-1_ Solyc09g059430 [no stop] | 10346 | Entry clone for C-terminal Myc-tag fusion | This work |
| pJLSmart | S/Rin4-2_ Solyc06g083390 | 10352 | Entry clone for N-terminal Myc-tag fusion | This work |
| pJLSmart | S/Rin4-2_ Solyc06g083390 [no stop] | 10348 | Entry clone for C-terminal Myc-tag fusion | This work |
| pJLSmart | S/Rin4-3_ Solyc12g098440 | 10350 | Entry clone for N-terminal Myc-tag fusion | This work |
| pJLSmart | S/Rin4-3_ Solyc12g098440 [no stop] | 10349 | Entry clone for C-terminal Myc-tag fusion | This work |
| pGWB417 | - | - | Binary Gateway destination vector for plant expression with C-terminal Myc-tag. | Nakagawa *et al.* (2007) |
| pGWB518 | - | - | Binary Gateway destination vector for plant expression with C-terminal Myc-tag. | Nakagawa *et al.* (2007) |
| pGWB518 | S/Rin4-1_ Solyc09g059430 | pKS13 | Generating N-terminal Myc-tag fusion | This work |
| pGWB518 | S/Rin4-2_ Solyc06g083390 | pKS15 | Generating N-terminal Myc-tag fusion | This work |
| pGWB518 | S/Rin4-3_ Solyc12g098440 | pKS17 | Generating N-terminal Myc-tag fusion | This work |
| pGWB518 | AtRIN4_At3G25070.1 | pKS18 | Generating N-terminal Myc-tag fusion | This work |
| pGWB417 | S/Rin4-1_ Solyc09g059430 | pKS12 | Generating C-terminal Myc-tag fusion | This work |
| pGWB417 | S/Rin4-2_ Solyc06g083390 | pKS14 | Generating C-terminal Myc-tag fusion | This work |
| pGWB417 | S/Rin4-3_ Solyc12g098440 | pKS16 | Generating C-terminal Myc-tag fusion | This work |
| pGWB417 | *avrRpt2_Pseudomonas syringae* pv. tomato NY15125 | pCM1 | Generating C-terminal Myc-tag fusion | Mazo-Molina *et al.* (2019) |
| pGWB417 | AvrRpt2(C122A) | - | Generating C-terminal Myc-tag fusion | Mazo-Molina *et al.* (2019) |
| pGWB417 | RipBN | - | Generating C-terminal Myc-tag fusion | Mazo-Molina *et al.* (2019) |
| pGWB417 | YFP | - | Generating C-terminal Myc-tag fusion | Roberts *et al.* (2019) |
| pBTEx | Prf(D1416V) | - | Generating N-terminal HA-tag fusion | Du *et al.* (2012) |
| pDONR/Zeo | - | - | Binary Gateway entry vector | Invitrogen |
| pDONR/Zeo | Ptr1-TRV2 | pSM1 | Entry clone for Ptr1-TRV2 construct | This work |
| pQ11 | TRV1 | - | VIGS TRV1 construct | Liu *et al.* (2002) |
| pQ11 | - | - | empty TRV2 construct | Liu *et al.* (2002) |
| pQ11 | EC1-TRV2 | - | VIGS TRV2 construct to silence EC1 | Rosli *et al.* (2013) |
| pQ11 | Ptr1-TRV2 | pSM2 | VIGS TRV2 construct to silence Ptr1 | This work |
| pBTEx | synPtr1 | pSM3 | Generating N-terminal HA-tag fusion | This work |
| pBTEx | StPtr1/RGA1 | pCM68 | Generating N-terminal HA-tag fusion | This work |

**Figure S1.**
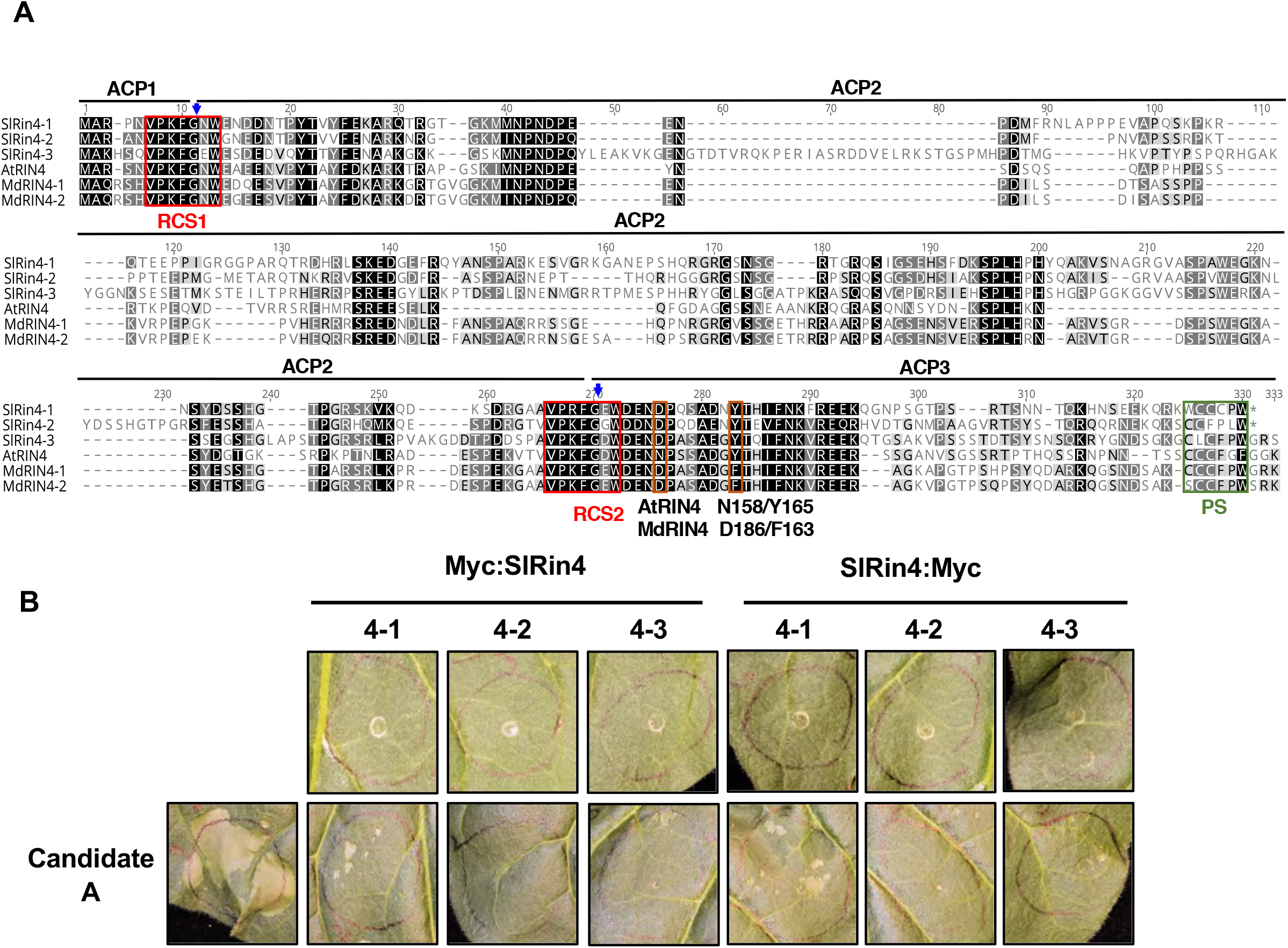
Amino acid sequences of Rin4 proteins in tomato and *Arabidopsis* and suppression of candidate A-mediated cell death by N- or C-terminal Myc-tagged tomato Rin4 proteins. **A)** Amino acid sequence alignment of Rin4 proteins from tomato and AtRIN4 from *Arabidopsis*. Black background indicates identical amino acid residues; grey background indicate similar amino acid residues; red boxes indicate regions (RCS1 and RCS2) cleaved by AvrRpt2 cysteine protease activity with blue arrows marking the exact cleavage site. AvrRpt2 cleavage products (ACP) are indicated. Green box indicates the palmitoylation site (PS). The polymorphic residues in AtRIN4 and the two MdRIN4 proteins are shown. **B)** *N. glutinosa* leaves were syringe-infiltrated with *Agrobacterium* strains carrying *A:HA* (OD_600_ = 0.1) and Sl*Rin4:Myc* (OD_600_ = 0.2) with both being expressed by the CaMV 35S promoter. Tomato Rin4 proteins were c-Myc tagged at the N- or C-terminal as indicated. Photographs were taken 48 hr after agroinfiltration. Data shown are representative of three independent experiments, using three biological replicates and infiltrating two leaves per plant.

**Figure S2.**
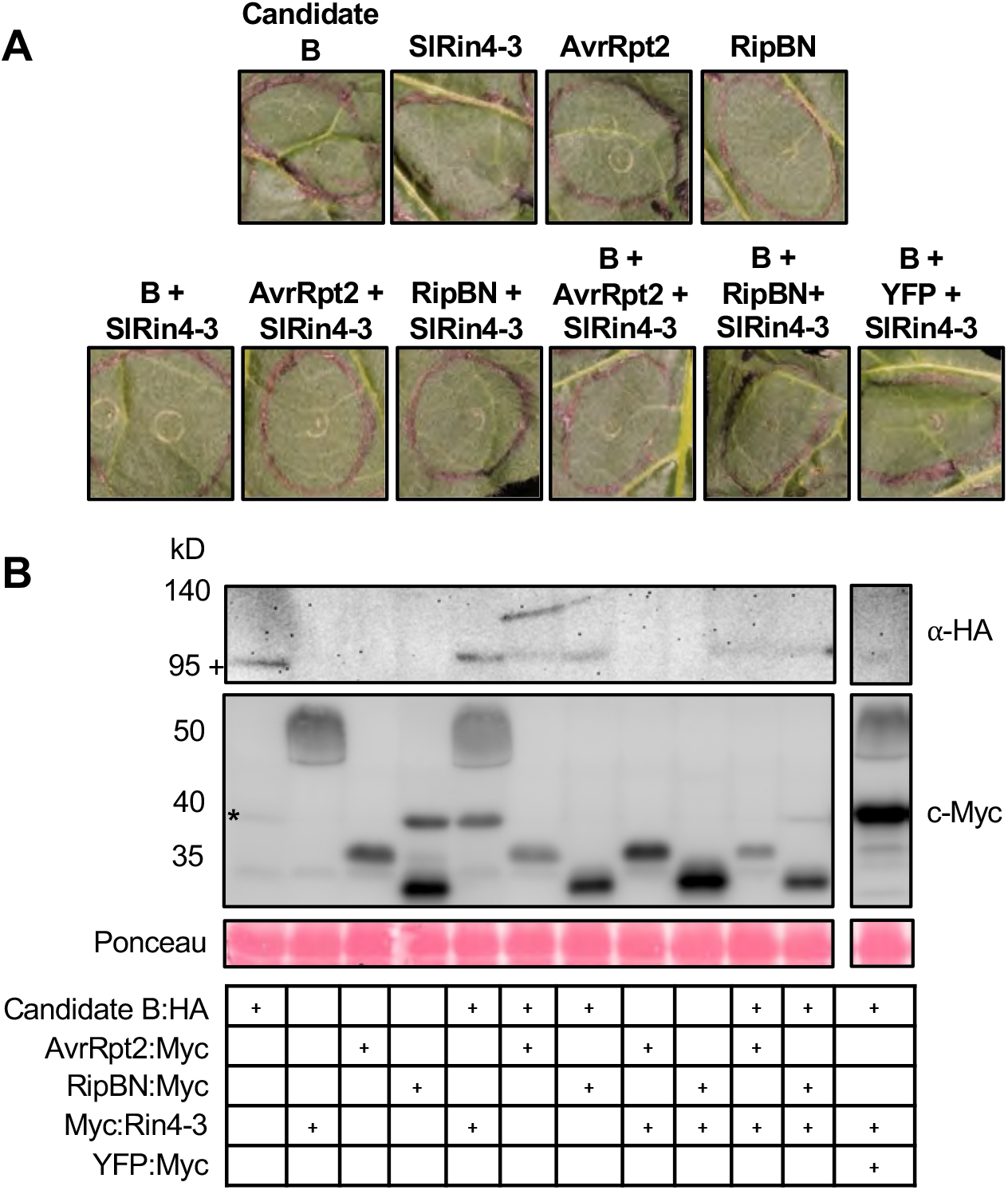
Candidate B does not mediate recognition of AvrRpt2 or RipBN in *N. glutinosa*. **A)** *N. glutinosa* leaves were syringe-infiltrated with *Agrobacterium* strains carrying either *B:HA* (OD_600_ = 0.1), *AvrRpt2:Myc* (OD_600_ = 0.05), *RipBN:Myc* (OD_600_ = 0.05), *YFP:Myc* (OD_600_ = 0.05) or *Myc*:Sl*Rin4-3* (OD_600_ = 0.2). Photographs were taken 48 hr after agroinfiltration. **B)** Immunoblot analysis of protein extracts isolated from *N. benthamiana* leaves agroinfiltrated with *B:HA* (OD_600_ = 0.1), *Myc*:Sl*Rin4-3* (OD_600_ = 0.2), *AvrRpt2:Myc* (OD_600_ = 0.05) or *YFP:Myc* (OD_600_ = 0.05). Samples were collected 28 hr after agroinfiltration. Total proteins extracted from infiltrated leaves were subjected to immunoblotting using an ∝-HA antibody to detect candidate B or a c-Myc antibody to detect *Sl*Rin4-3, AvrRpt2, RipBN and YFP. Protein masses are indicated at the left of the blot. Ponceau staining shows amount of protein loaded in each lane. +, candidate B protein; *, a nonspecific bands across the immunoblot. All genes in this figure were expressed from the CaMV 35S promoter. Data shown are representative of three independent experiments, using three biological replicates and infiltrating two leaves per plant.

**Figure S3.**
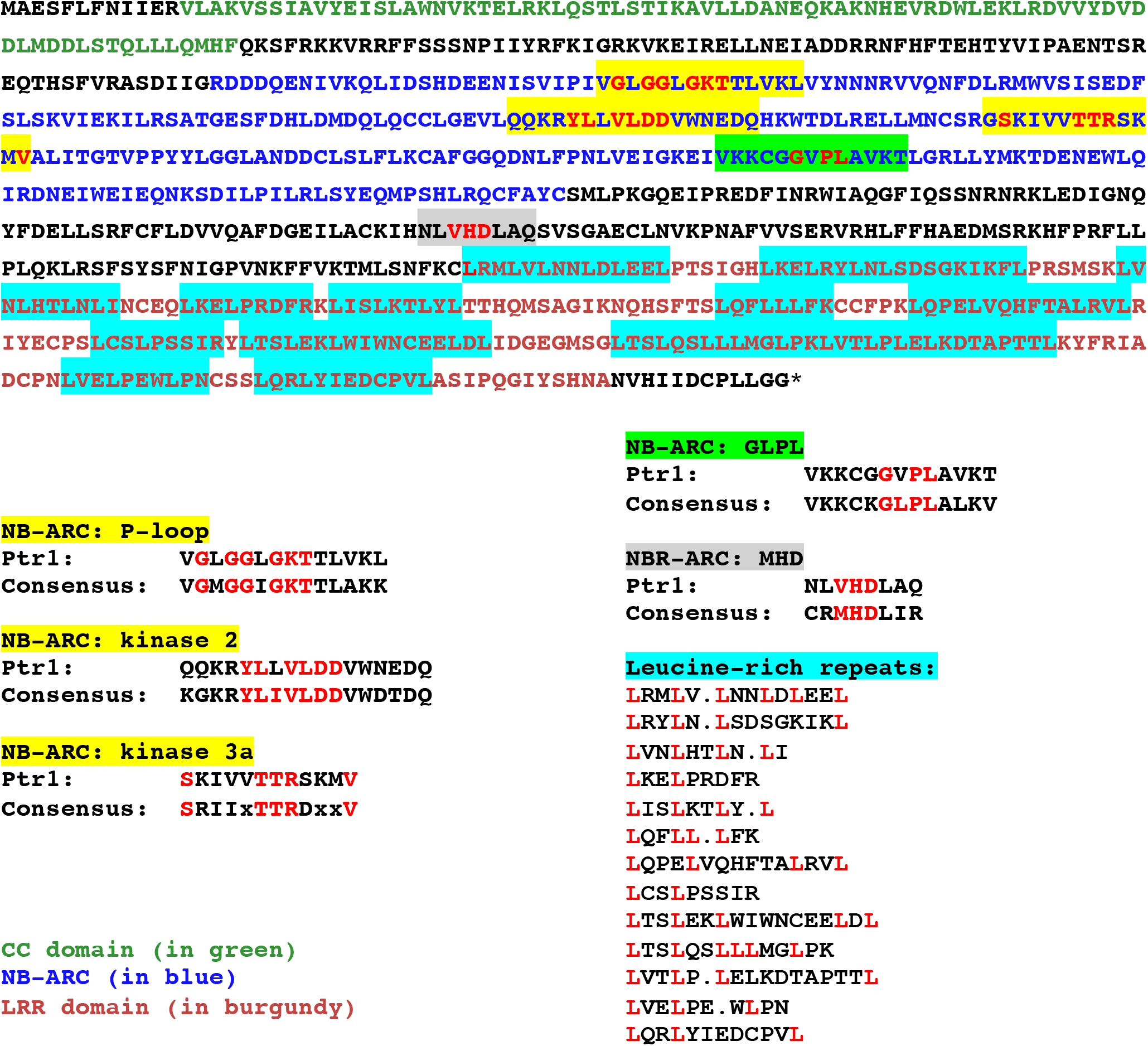
Amino acid sequence of the predicted Ptr1 protein showing motifs characteristic of NLR immune receptors. The LRR repeats (LxxLxxLxxLxLxxLxx motif) were identified using the LRRSearch Tool (Bej, et al., 2014) and adjusted manually.

**Figure S4.**
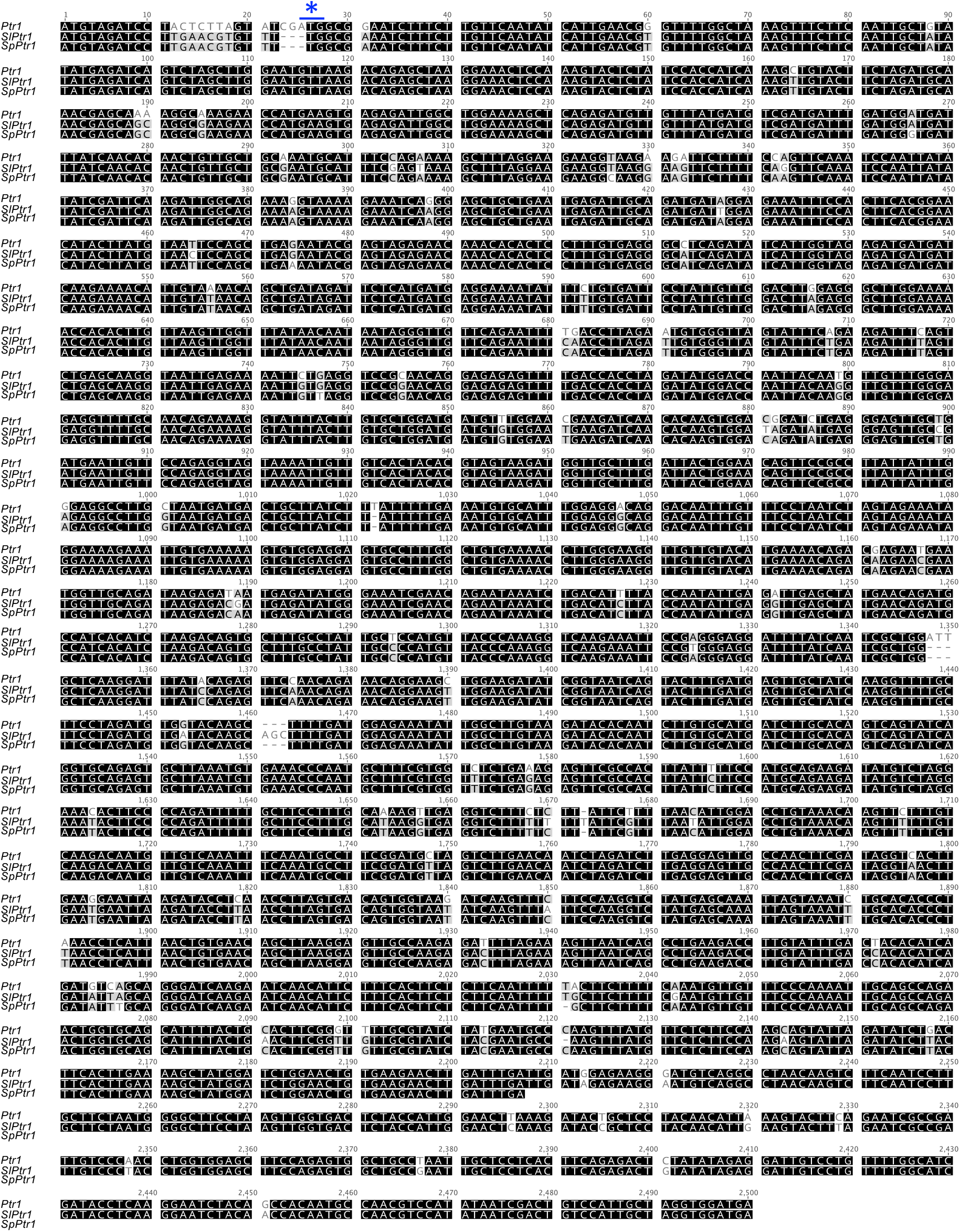
The ortholog of *Ptr1* is a pseudogene in tomato and *S. pennellii*. ClustalW alignment of the nucleotide sequence (including 20 nucleotides upstream of start codon) of candidate A and its ortholog in *S. lycopersicum* (*SlPtr1*) and *S. pennellii* (*SpPtr1*). The blue asterisk denotes the start codon unique to *S. lycopersicoides Ptr1*. Numbers on right correspond to the sequence of *Ptr1*.

**Figure S5.**
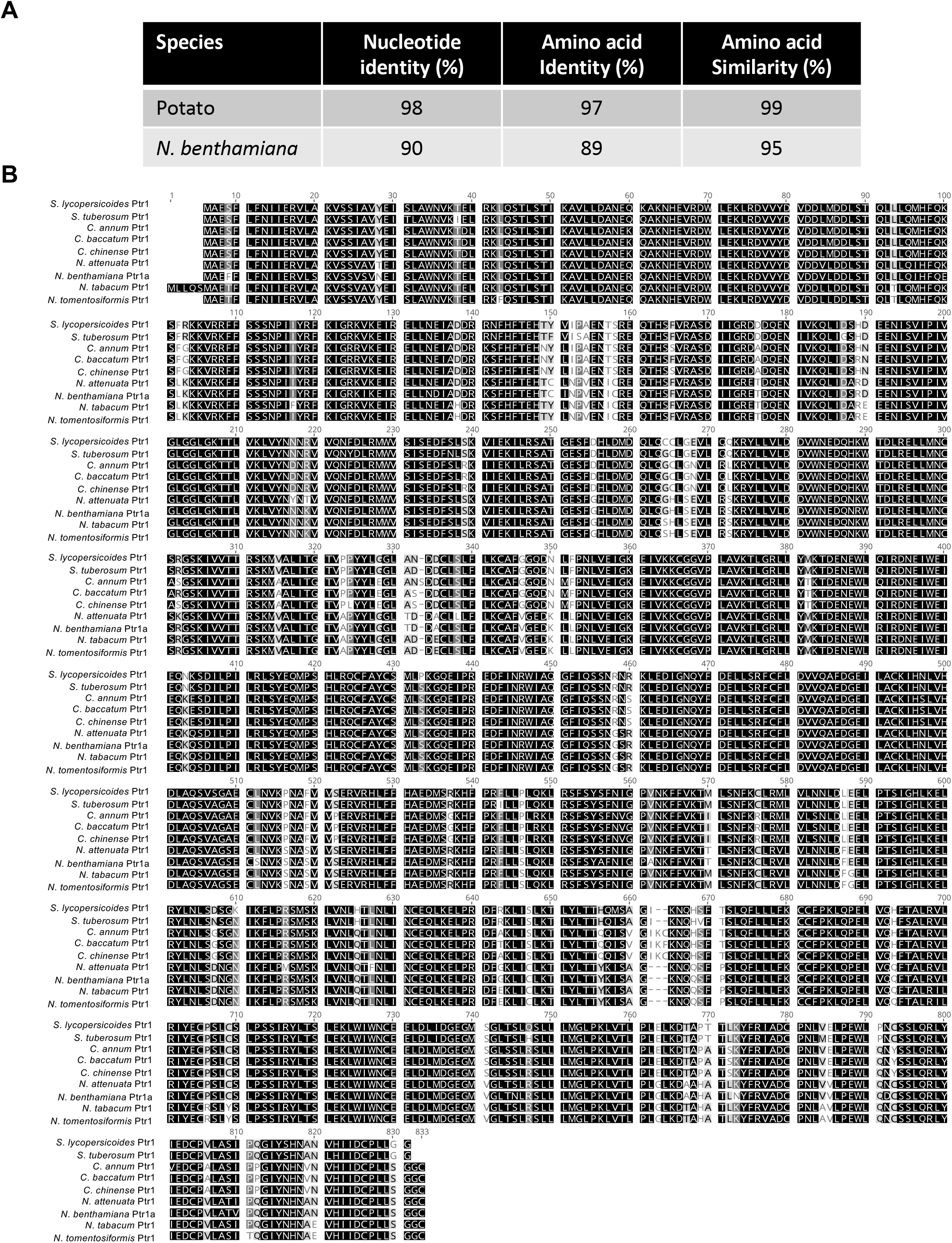
*Ptr1* orthologs in various Solanaceae species. **A)** Comparison of *Ptr1* nucleotide and amino acid sequences with its orthologs in potato and *Nicotiana benthamiana*. **B)** Amino acid sequence alignment of Ptr1 proteins from various Solanaceae species.

**Figure S6.**
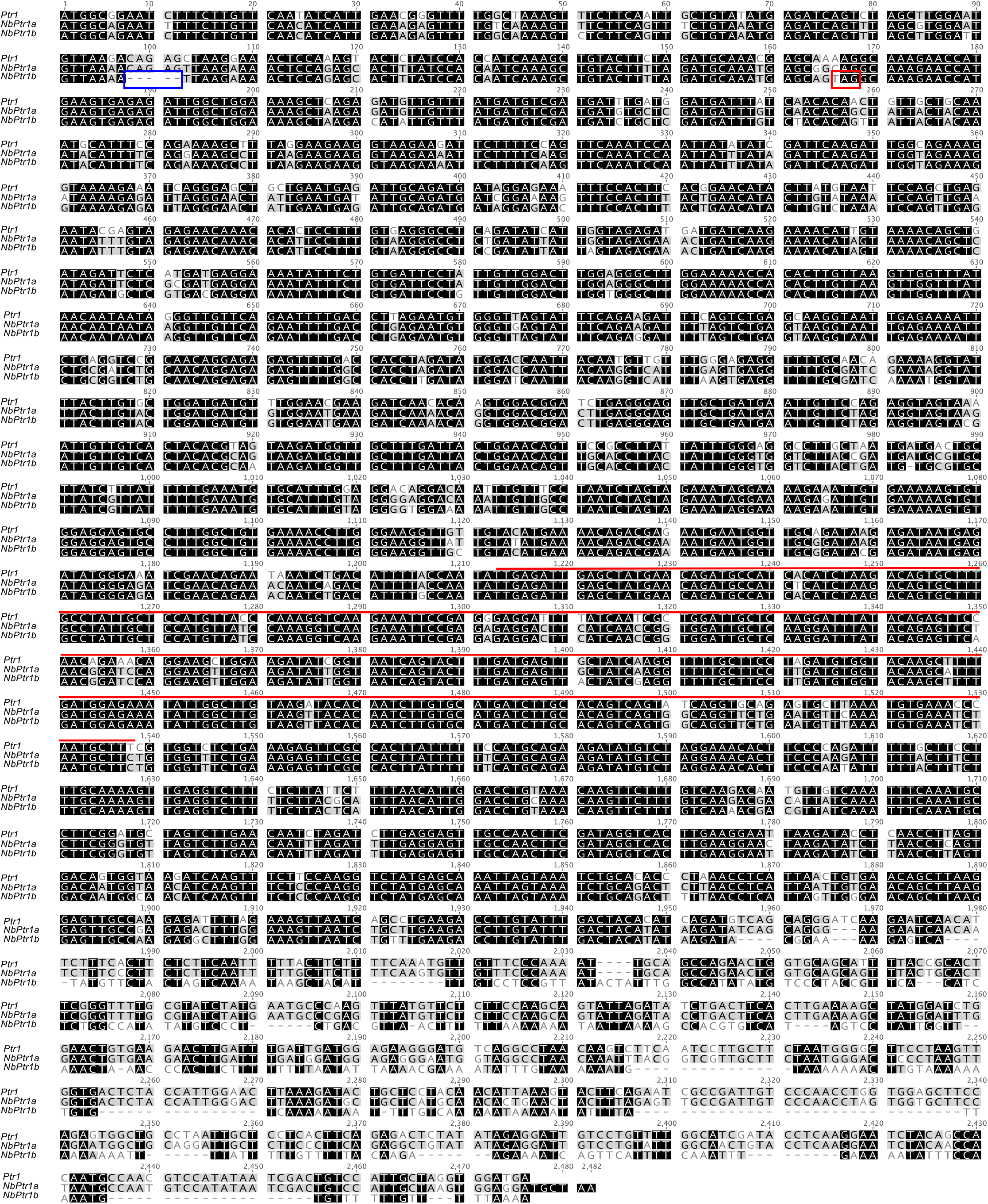
*Ptr1* has extensive sequence similarity with the two *Ptr1* homologs in *N. benthamiana*. Nucleotide alignment of *Ptr1*, *NbPtr1a* and *NbPtr1b*. A five base pair deletion (blue box) in *NbPtr1b* leads to a premature stop codon (red box), although the nucleotide sequence remains conserved at the region targeted by VIGS (red line).

**Figure S7.**
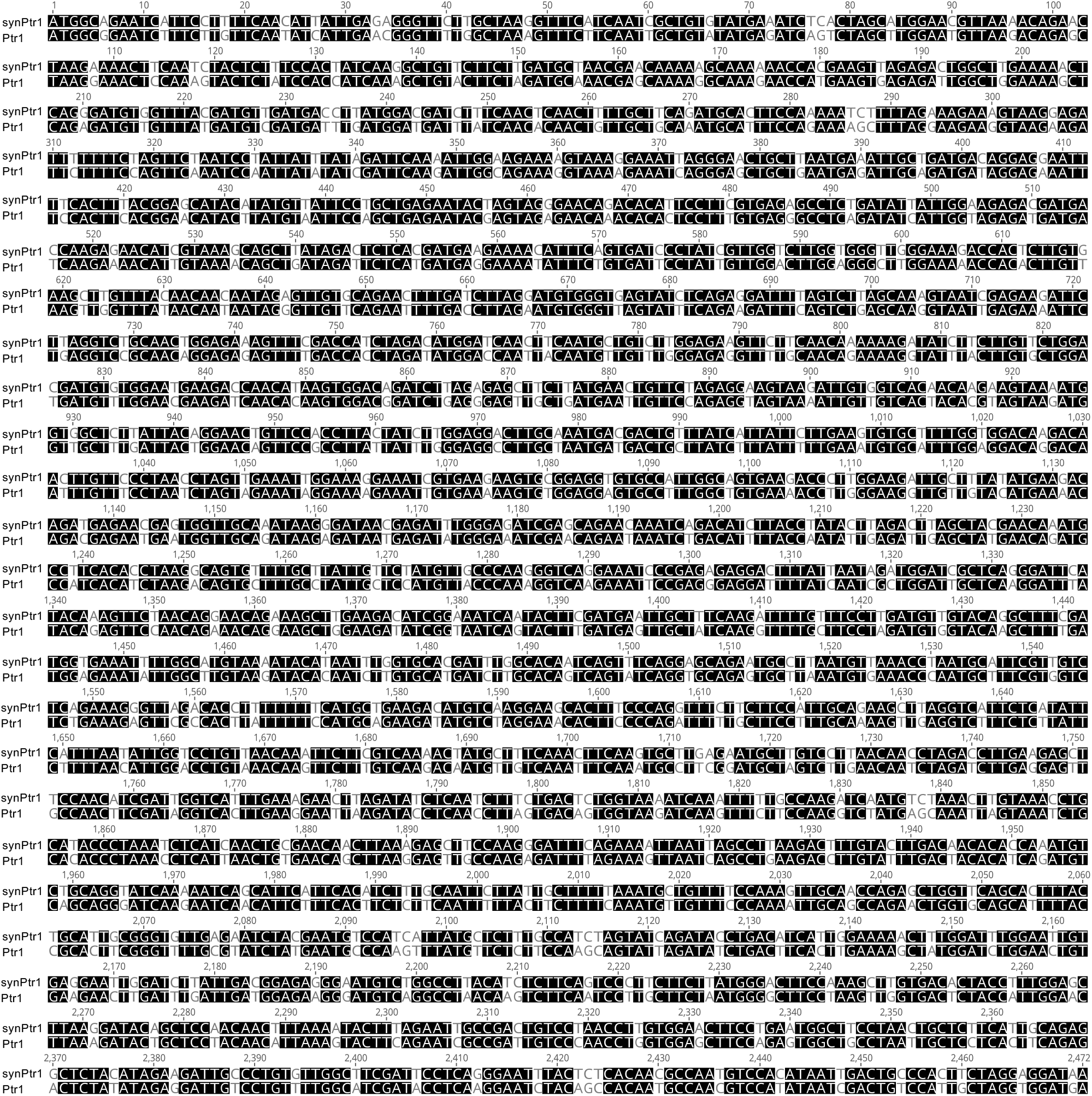
Nucleotide alignment of *Ptr1* and synthetic *Ptr1* (synPtr1). Identical nucleotides are shown in black. The amino acid sequence is identical for both Ptr1 and synPtr1.

**Figure S8.**
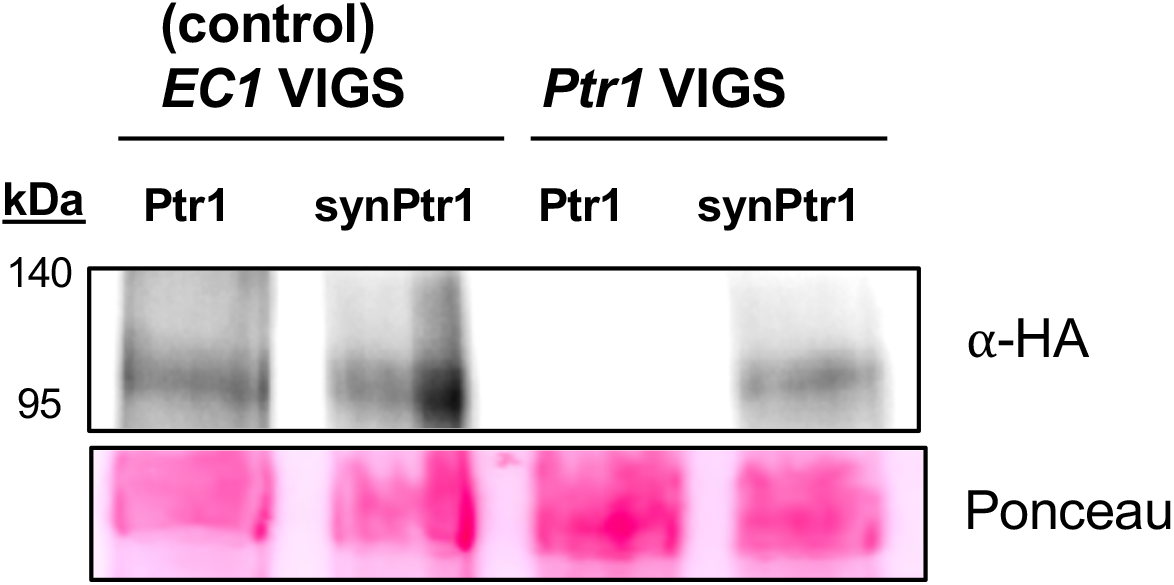
Immunoblot analysis of protein extracts isolated from *N. benthamiana* leaves transiently expressing *35S:Ptr1:HA* or *35S:synPtr1:HA*. Proteins were extracted from leaves collected 46 hr after *Agrobacterium*-mediated transient transformation of Ptr1 and synPtr1 (agroinfiltration; OD_600_ = 0.1) and subjected to immunoblotting using an ∝-HA antibody to detect Ptr1 and synPtr1. Protein masses are indicated at the left of each blot. Ponceau staining shows amount of protein loaded in each lane.

**Figure S9A.**
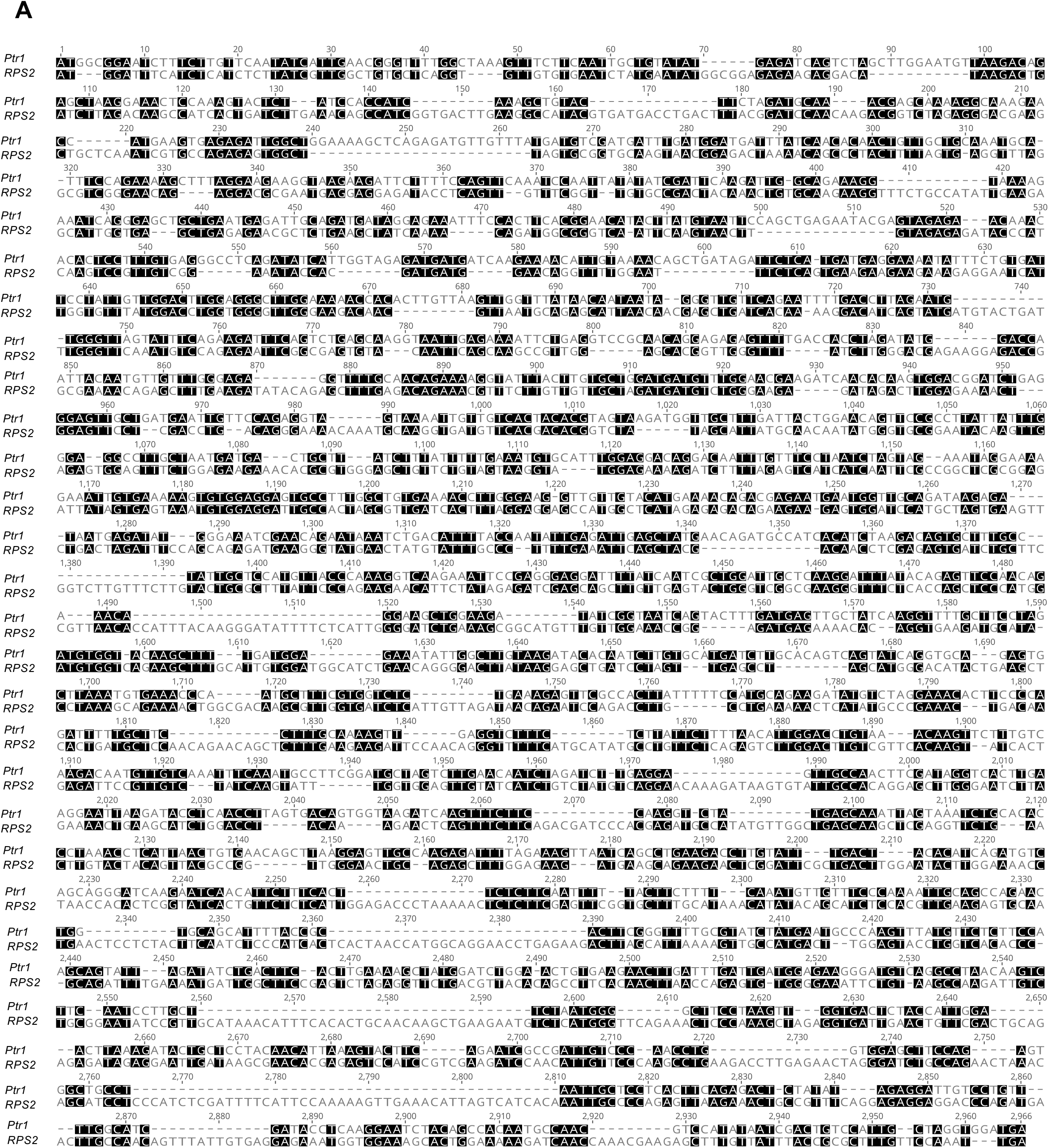
*Ptr1* has little nucleotide sequence similarity with *RPS2*. Alignment of the nucleotide sequences of Ptr1 and RPS2 was generated using MUSCLE. Identical nucleotides are shown in black.

**Figure S9B.**
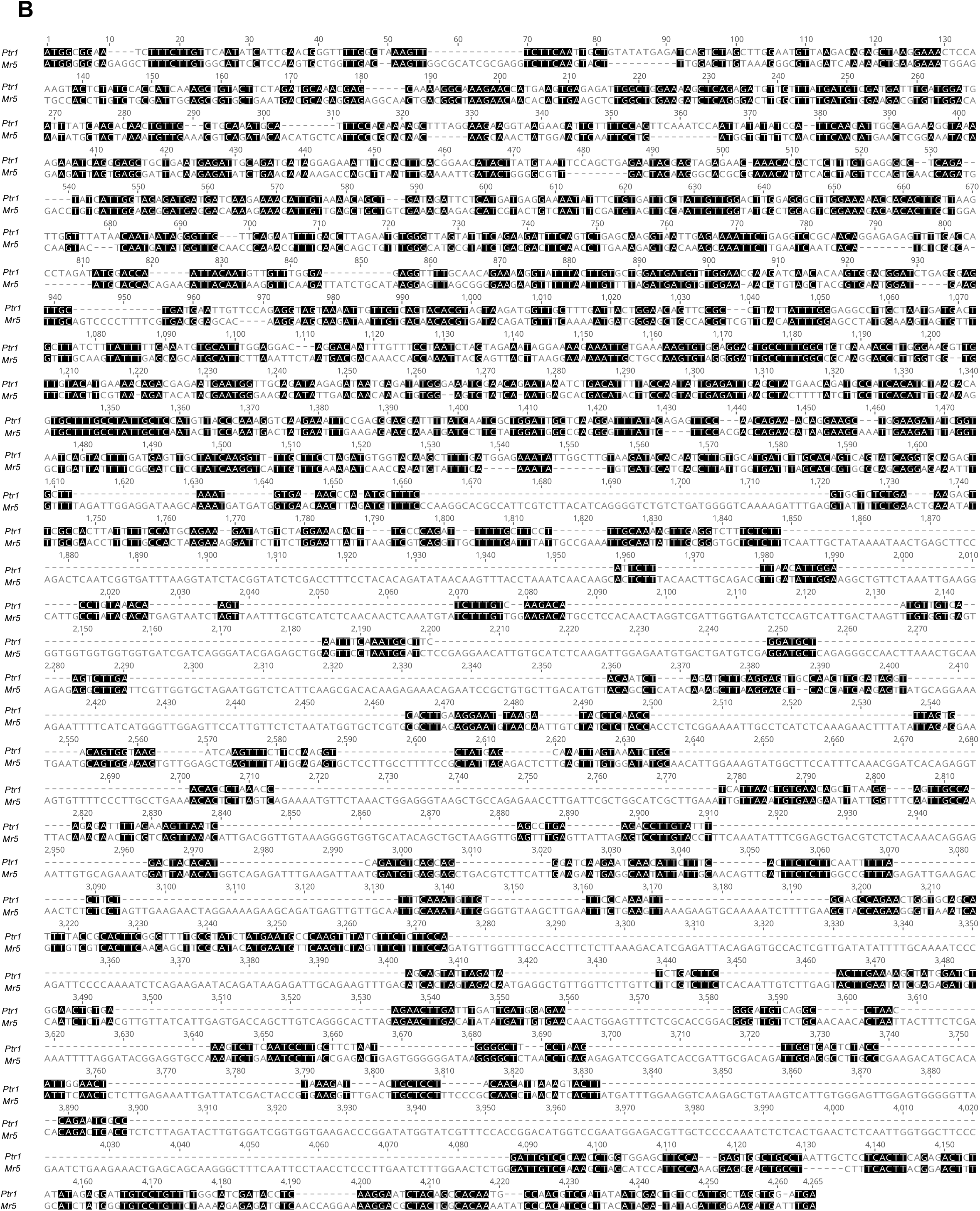
*Ptr1* has little nucleotide sequence similarity with *Mr5*. Alignment of the nucleotide sequences of Ptr1 and RPS2 was generated using MUSCLE. Identical nucleotides are shown in black.

**Figure S10A.**
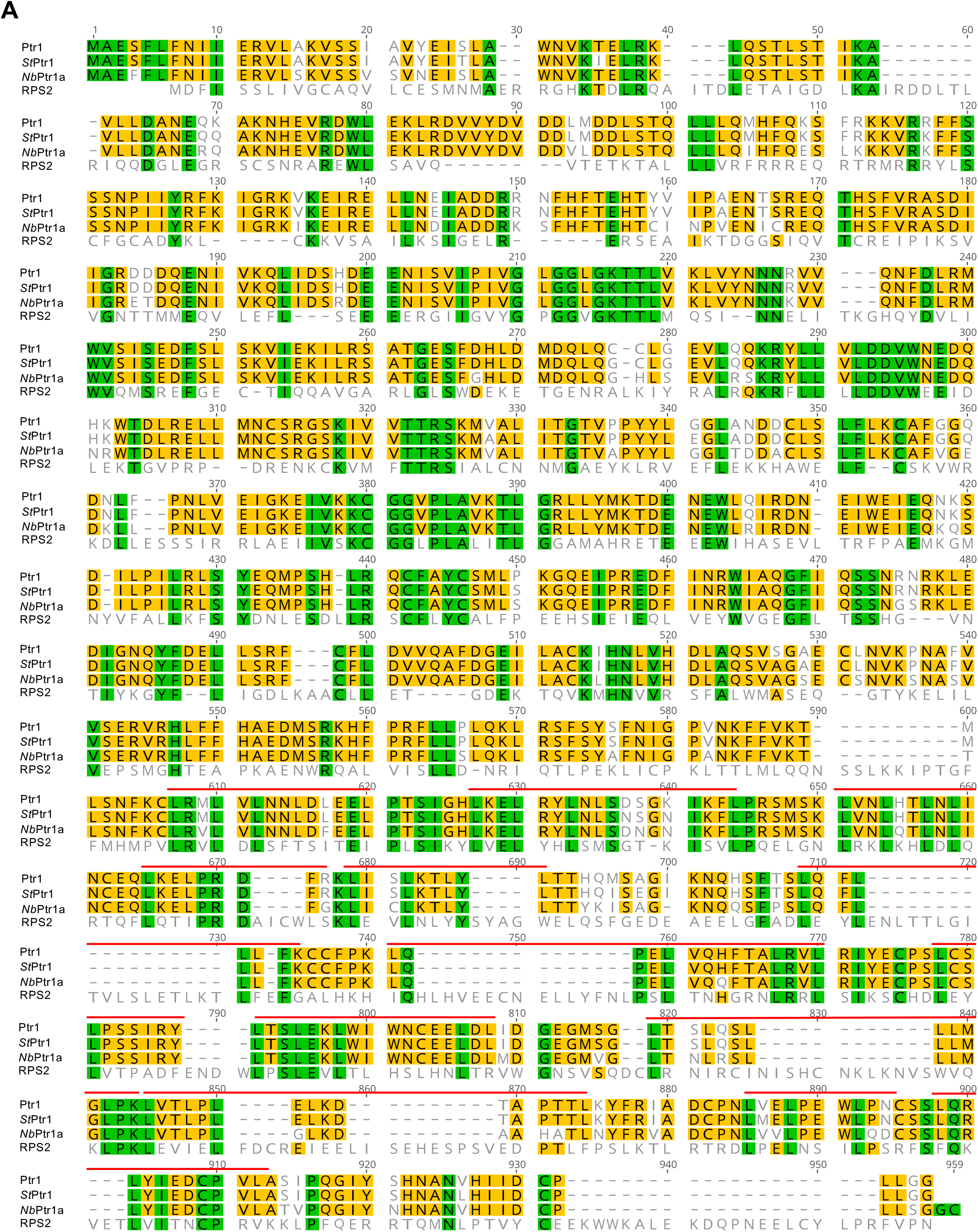
Ptr1, StPtr1 and NbPtr1 share little amino acid sequence similarity with RPS2. **A)** Amino acid alignment of the Ptr1 protein with StPtr1, NbPtr1a, and RPS2. Green indicates all four of the proteins have the same residue at that position, gold indicates that 3 of the 4 proteins have the same residue at that position. Red lines indicates leucine-rich repeats.

**Figure S10B.**
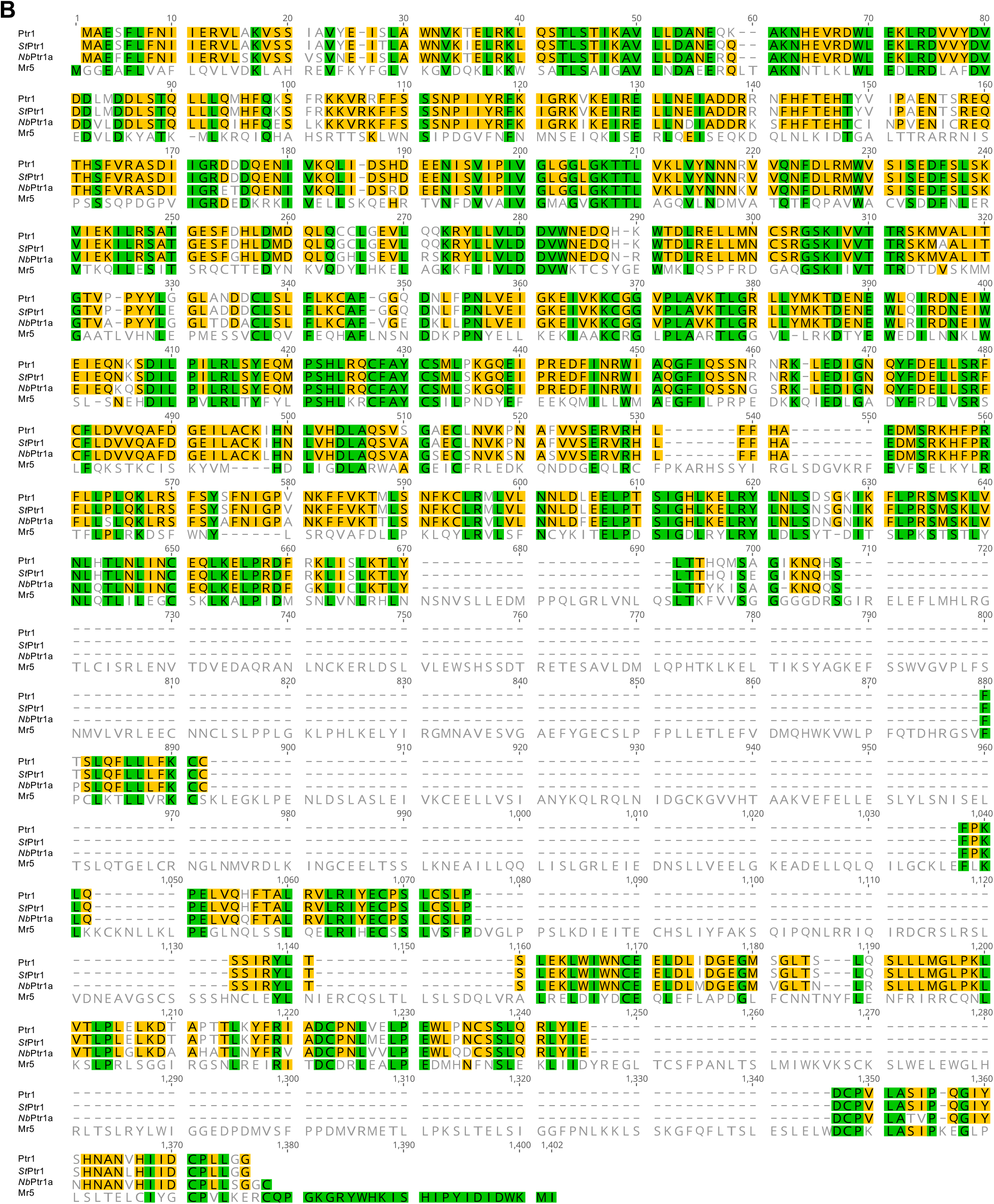
Ptr1, StPtr1 and NbPtr1 share little amino acid sequence similarity with Mr5. **B)** Amino acid alignment of the Ptr1 protein with StPtr1, NbPtr1a, and Mr5. Green indicates all four of the proteins have the same residue at that position, gold indicates that 3 of the 4 proteins have the same residue at that position.

**Figure S11.**
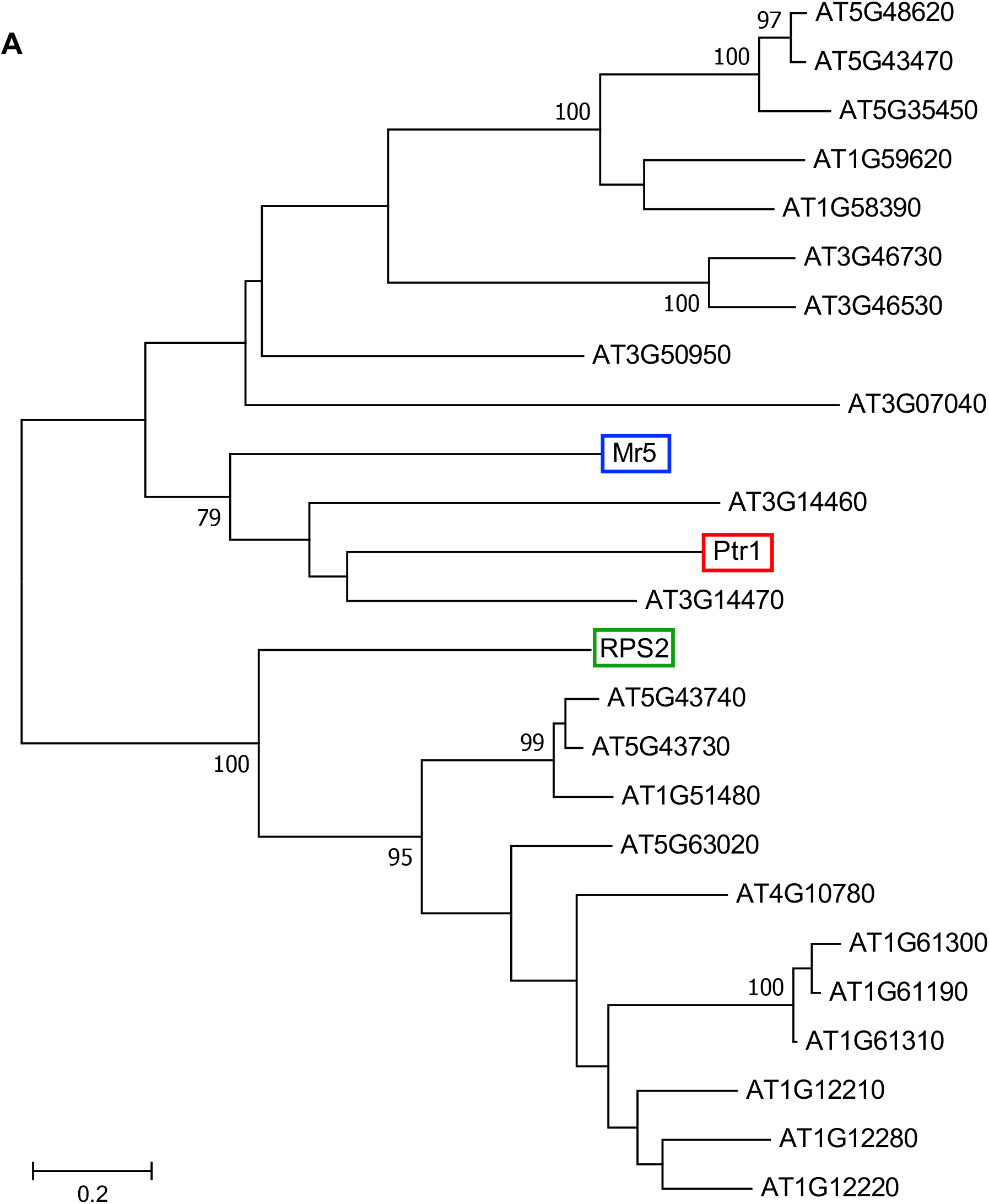
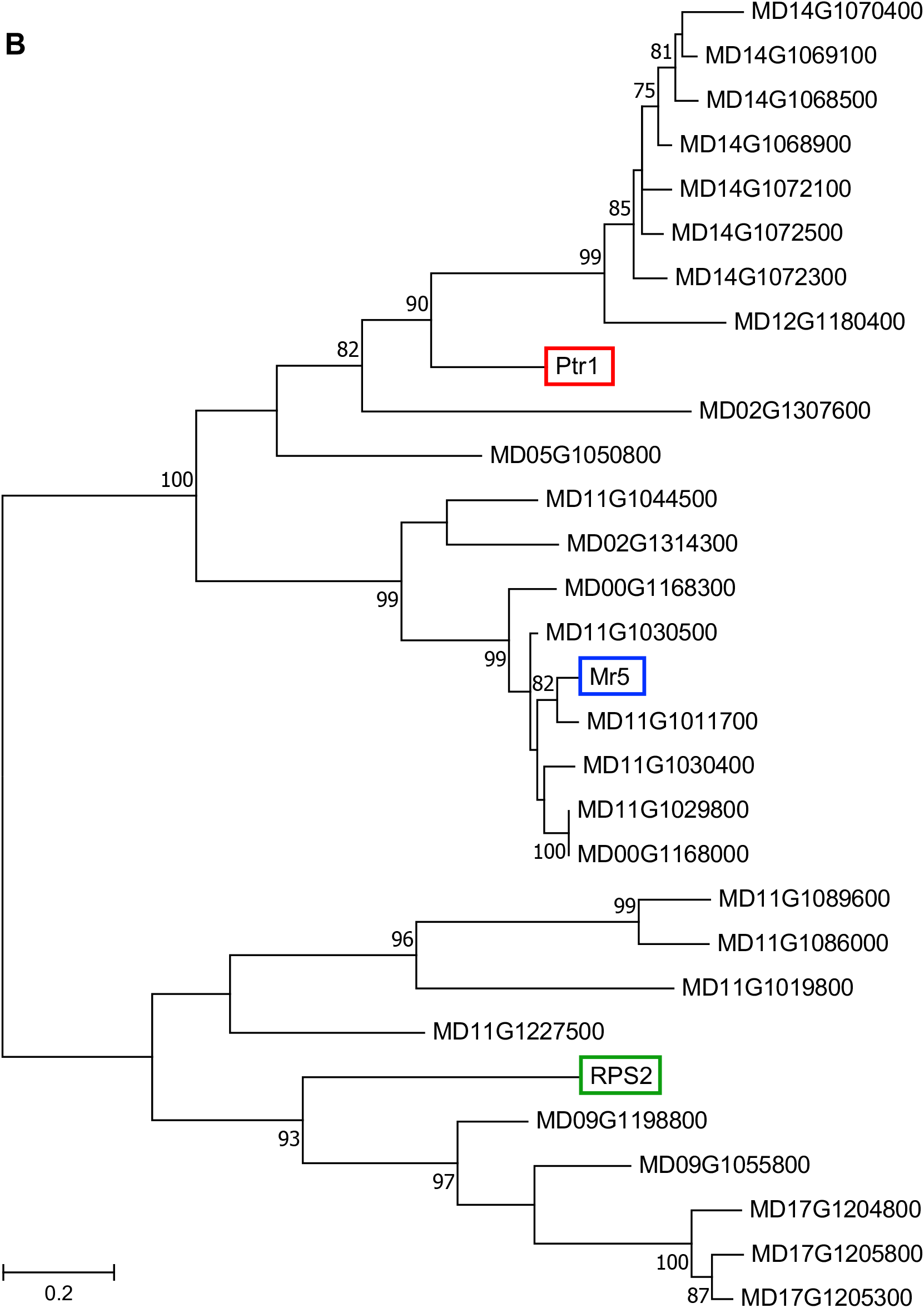
Phylogenetic tree of Ptr1, Mr5, RPS2 with their most closely related proteins in Arabidopsis and apple. Maximum likelihood tree of top BLAST hits of Ptr1, Mr5, and RPS2 NB-ARC domains from: **A)** Arabidopsis (AT; Araport 11 protein database) and **B)** *Malus domestica* (MD; GDDH13 V1.1 protein database). Only hits with an intact NB-ARC domain, including the P-loop, kinase 2, kinase 3a, and GLPL motifs were included in the analysis. Bootstrapping 1000 replicates, with bootstraps over 75 shown. The tree is drawn to scale, with branch lengths measured in the number of substitutions per site. See next page for part B.

**Figure S12.**
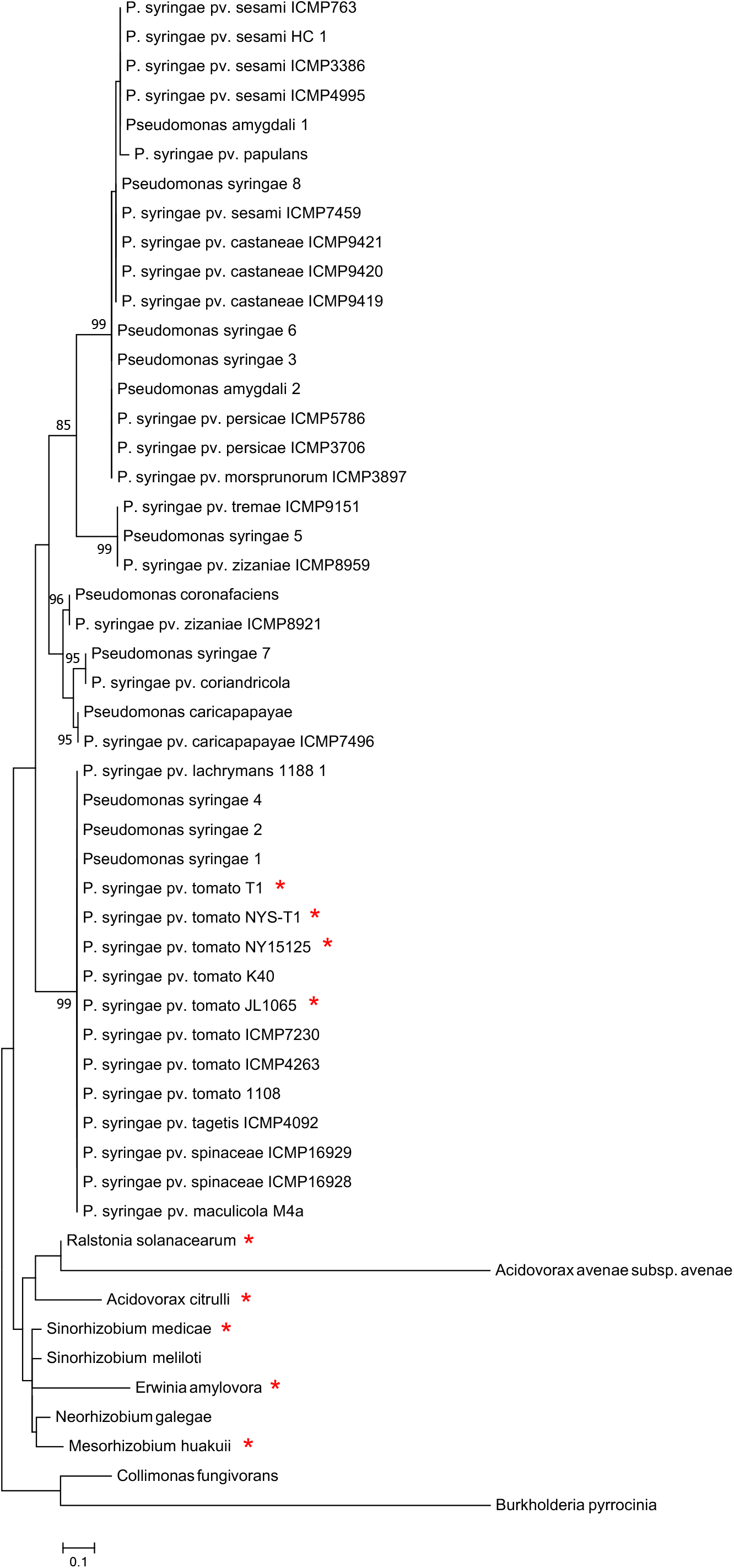
Maximum likelihood phylogenetic tree of AvrRpt2 homologs in Genbank. Ptr1 is known to mediate recognition of AvrRpt2 proteins indicated with a red asterisk (all are cysteine protease-active). Bootstrapping 1000 replicates, with bootstraps over 75 shown. The tree is drawn to scale, with branch lengths measured in the number of substitutions per site.

## References

Anzalone, A.V., Randolph, P.B., Davis, J.R., Sousa, A.A., Koblan, L.W., Levy, J.M., Chen, P.J., Wilson, C., Newby, G.A., Raguram, A. and Liu, D.R. (2019) Search-and-replace genome editing without double-strand breaks or donor DNA. Nature, 576, 149–157.

Ashfield, T., Ong, L.E., Nobuta, K., Schneider, C.M. and Innes, R.W. (2004) Convergent evolution of disease resistance gene specificity in two flowering plant families. Plant Cell, 16, 309–318.

Axtell, M.J., Chisholm, S.T., Dahlbeck, D. and Staskawicz, B.J. (2003) Genetic and molecular evidence that the *Pseudomonas syringae* type III effector protein AvrRpt2 is a cysteine protease. Mol Microbiol, 49, 1537–1546.

Axtell, M.J. and Staskawicz, B.J. (2003) Initiation of RPS2-specified disease resistance in *Arabidopsis* is coupled to the AvrRpt2-directed elimination of RIN4. Cell, 112, 369–377.

Bao, Z., Meng, F., Strickler, S.R., Dunham, D.M., Munkvold, K.R. and Martin, G.B. (2015) Identification of a candidate gene in *Solanum habrochaites* for resistance to a race 1 strain of *Pseudomonas syringae* pv. tomato. The Plant Genome, 8, 10.3835/plantgenome2015.3802.0006.

Bernatzky, R. and Tanksley, S.D. (1986) Genetics of actin-related sequences in tomato. Theor Appl Genet, 72, 314–321.

Bolger, A., Scossa, F., Bolger, M.E., Lanz, C., Maumus, F., Tohge, T., Quesneville, H., Alseekh, S., Sorensen, I., Lichtenstein, G., Fich, E.A., Conte, M., Keller, H., Schneeberger, K., Schwacke, R., Ofner, I., Vrebalov, J., Xu, Y., Osorio, S., Aflitos, S.A., Schijlen, E., Jimenez-Gomez, J.M., Ryngajllo, M., Kimura, S., Kumar, R., Koenig, D., Headland, L.R., Maloof, J.N., Sinha, N., van Ham, R.C., Lankhorst, R.K., Mao, L., Vogel, A., Arsova, B., Panstruga, R., Fei, Z., Rose, J.K., Zamir, D., Carrari, F., Giovannoni, J.J., Weigel, D., Usadel, B. and Fernie, A.R. (2014a) The genome of the stress-tolerant wild tomato species *Solanum pennellii*. Nat Genet, 46, 1034–1038.

Bolger, A.M., Lohse, M. and Usadel, B. (2014b) Trimmomatic: A flexible trimmer for Illumina sequence data. Bioinformatics, 30, 2114–2120.

Bombarely, A., Rosli, H.G., Vrebalov, J., Moffett, P., Mueller, L.A. and Martin, G.B. (2012) A draft genome sequence of *Nicotiana benthamiana* to enhance molecular plant-microbe biology research. Mol Plant-Microbe Interact, 25, 1523–1530.

Canady, M.A., Meglic, V. and Chetelat, R.T. (2005) A library of *Solanum lycopersicoides* introgression lines in cultivated tomato. Genome, 48, 685–697.

Carter, M.E., Helm, M., Chapman, A., Wan, E., Restrepo Sierra, A.M., Innes, R., Bogdanove, A.J. and Wise, R.P. (2018) Convergent evolution of effector protease recognition by Arabidopsis and barley. Mol Plant Microbe Interact, 32, 550–565.

Chakravarthy, S., Velasquez, A.C., Ekengren, S.K., Collmer, A. and Martin, G.B. (2010) Identification of *Nicotiana benthamiana* genes involved in pathogen-associated molecular pattern-triggered immunity. Mol Plant Microbe Interact, 23, 715–726.

Coaker, G., Falick, A. and Staskawicz, B. (2005) Activation of a phytopathogenic bacterial effector protein by a eukaryotic cyclophilin. Science, 308, 548–550.

Day, B., Dahlbeck, D., Huang, J., Chisholm, S.T., Li, D. and Staskawicz, B.J. (2005) Molecular basis for the RIN4 negative regulation of RPS2 disease resistance. Plant Cell, 17, 1292–1305.

de la Torre, F., Gutierrez-Beltran, E., Pareja-Jaime, Y., Chakravarthy, S., Martin, G.B. and del Pozo, O. (2013) The tomato calcium sensor Cbl10 and its interacting protein kinase Cipk6 define a signaling pathway in plant immunity. Plant Cell, 25, 2748–2764.

del Pozo, O., Pedley, K.F. and Martin, G.B. (2004) MAPKKKα is a positive regulator of cell death associated with both plant immunity and disease. EMBO J, 23, 3072–3082.

Delcher, A.L., Kasif, S., Fleischmann, R.D., Peterson, J., White, O. and Salzberg, S.L. (1999) Alignment of whole genomes. Nucleic Acids Res, 27, 2369–2376.

Dillon, M.M., Almeida, R.N.D., Laflamme, B., Martel, A., Weir, B.S., Desveaux, D. and Guttman, D.S. (2019) Molecular evolution of *Pseudomonas syringae* type III secreted effector proteins. Front Plant Sci, 10, 418.

Dobin, A., Davis, C.A., Schlesinger, F., Drenkow, J., Zaleski, C., Jha, S., Batut, P., Chaisson, M. and Gingeras, T.R. (2013) STAR: ultrafast universal RNA-seq aligner. Bioinformatics, 29, 15–21.

Ekengren, S., Liu, Y., Schiff, M., Dinesh-Kumar, S. and Martin, G. (2003) Two MAPK cascades, NPR1, and TGA transcription factors play a role in Pto-mediated disease resistance in tomato. Plant J, 36, 905–917.

Eschen-Lippold, L., Jiang, X., Elmore, J.M., Mackey, D.M., Shan, L., Coaker, G.L., Scheel, D. and Lee, J. (2016) Bacterial AvrRpt2-like cysteine proteases block activation of the Arabidopsis mitogen-activated protein kinases, MPK4 and MPK11. Plant Physiol, 171, 2223–2238.

Fahrentrapp, J., Broggini, G.A.L., Kellerhals, M., Peil, A., Richter, K., Zini, E. and Gessler, C. (2012) A candidate gene for fire blight resistance in Malus × robusta 5 is coding for a CC–NBS–LRR. Tree Genetics & Genomics, 9, 237–251.

Felsenstein, J. (1985) Confidence limits on phylogenies: An approach using the bootstrap. Evolution, 39, 783–791.

Fernandez-Pozo, N., Rosli, H.G., Martin, G.B. and Mueller, L.A. (2015) The SGN VIGS Tool: User-friendly software to design virus-induced gene silencing (VIGS) constructs for functional genomics. Mol Plant, 8, 486–488.

Gendrel, A.V., Lippman, Z., Martienssen, R. and Colot, V. (2005) Profiling histone modification patterns in plants using genomic tiling microarrays. Nat Methods, 2, 213–218.

Hassan, J.A., Zhou, Y.J. and Lewis, J.D. (2017) A rapid seedling resistance assay identifies wild tomato lines that are resistant to *Pseudomonas syringae* pv. tomato race 1. Mol Plant Microbe Interact, 30, 701–709.

Holsters, M., Silva, B., Van Vliet, F., Genetello, C., De Block, M., Dhaese, P., Depicker, A., Inze, D., Engler, G., Villarroel, R. and, et al. (1980) The functional organization of the nopaline *A. tumefaciens* plasmid pTiC58. Plasmid, 3, 212–230.

Jones, D.T., Taylor, W.R. and Thornton, J.M. (1992) The rapid generation of mutation data matrices from protein sequences. Comput Appl Biosci, 8, 275–282.

Jones, J.B. (1991) Bacterial speck. In Compendium of tomato diseases (Jones, J.B., Jones, J.P., Stall, R.E. and Zitter, T.A. eds). St. Paul, MN: APS Press, pp. 26–27.

Jung, S., Lee, T., Cheng, C.H., Buble, K., Zheng, P., Yu, J., Humann, J., Ficklin, S.P., Gasic, K., Scott, K., Frank, M., Ru, S., Hough, H., Evans, K., Peace, C., Olmstead, M., DeVetter, L.W., McFerson, J., Coe, M., Wegrzyn, J.L., Staton, M.E., Abbott, A.G. and Main, D. (2019) 15 years of GDR: New data and functionality in the Genome Database for Rosaceae. Nucleic Acids Res, 47, D1137–D1145.

Kessens, R., Ashfield, T., Kim, S.H. and Innes, R.W. (2014) Determining the GmRIN4 requirements of the soybean disease resistance proteins Rpg1b and Rpg1r using a *Nicotiana glutinosa*-based agroinfiltration system. PLoS One, 9, e108159.

Kim, H.S., Desveaux, D., Singer, A.U., Patel, P., Sondek, J. and Dangl, J.L. (2005) The *Pseudomonas syringae* effector AvrRpt2 cleaves its C-terminally acylated target, RIN4, from Arabidopsis membranes to block RPM1 activation. Proc Natl Acad Sci USA, 102, 6496–6501.

Kim, Y., Lin, N. and Martin, G. (2002) Two distinct *Pseudomonas* effector proteins interact with the Pto kinase and activate plant immunity. Cell, 109, 589–598.

Kraus, C.M., Mazo-Molina, C., Smart, C.D. and Martin, G.B. (2017) *Pseudomonas syringae* pv. tomato strains from New York exhibit virulence attributes intermediate between typical race 0 and race 1 strains. Plant Disease, 101, 1442–1448.

Kud, J., Zhao, Z., Du, X., Liu, Y., Zhao, Y. and Xiao, F. (2013) SGT1 interacts with the Prf resistance protein and is required for Prf accumulation and Prf-mediated defense signaling. Biochem Biophys Res Comm, 431, 501–505.

Kumar, S., Stecher, G. and Tamura, K. (2016) MEGA7: Molecular Evolutionary Genetics Analysis version 7.0 for bigger datasets. Mol Biol Evol, 33, 1870–1874.

Kunkeaw, S., Tan, S. and Coaker, G. (2010) Molecular and evolutionary analyses of *Pseudomonas syringae* pv. tomato race 1. Mol Plant-Microbe Interact, 23, 415–424.

Langmead, B. and Salzberg, S.L. (2012) Fast gapped-read alignment with Bowtie 2. Nat Methods, 9, 357–359.

Lim, M.T. and Kunkel, B.N. (2005) The *Pseudomonas syringae avrRpt2* gene contributes to virulence on tomato. Mol Plant Microbe Interact, 18, 626–633.

Liu, Y., Schiff, M. and Dinesh-Kumar, S.P. (2002) Virus-induced gene silencing in tomato. Plant J, 31, 777–786.

Lu, R., Malcuit, I., Moffett, P., Ruiz, M.T., Peart, J., Wu, A.J., Rathjen, J.P., Bendahmane, A., Day, L. and Baulcombe, D.C. (2003) High throughput virus-induced gene silencing implicates heat shock protein 90 in plant disease resistance. EMBO J, 22, 5690–5699.

Mackey, D., Belkhadir, Y., Alonso, J.M., Ecker, J.R. and Dangl, J.L. (2003) Arabidopsis RIN4 is a target of the type III virulence effector AvrRpt2 and modulates RPS2-mediated resistance. Cell, 112, 379–389.

Marcais, G., Delcher, A.L., Phillippy, A.M., Coston, R., Salzberg, S.L. and Zimin, A. (2018) MUMmer4: A fast and versatile genome alignment system. PLoS Comput Biol, 14, e1005944.

Martin, G.B. (2012) Suppression and activation of the plant immune system by Pseudomonas syringae effectors AvrPto and AvrPtoB. In Effectors in Plant-Microbe Interactions (Martin, F. and Kamoun, S. eds). Ames, IA: Wiley-Blackwell, pp. 123–154.

Mathieu, J., Schwizer, S. and Martin, G.B. (2014) Pto kinase binds two domains of AvrPtoB and its proximity to the effector E3 ligase determines if it evades degradation and activates plant immunity. PLoS Pathog, 10, e1004227.

Mazo-Molina, C., Mainiero, S., Hind, S.R., Kraus, C.M., Vachev, M., Maviane-Macia, F., Lindeberg, M., Saha, S., Strickler, S.R., Feder, A., Giovannoni, J.J., Smart, C.D., Peeters, N. and Martin, G.B. (2019) The *Ptr1* locus of *Solanum lycopersicoides* confers resistance to race 1 strains of *Pseudomonas syringae* pv. tomato and to *Ralstonia pseudosolanacearum* by recognizing the type III effectors AvrRpt2 and RipBN. Mol Plant Microbe Interact, 32, 949–960.

Mucyn, T.S., Wu, A.J., Balmuth, A.L., Arasteh, J.M. and Rathjen, J.P. (2009) Regulation of tomato Prf by Pto-like protein kinases. Mol Plant-Microbe Interact, 22, 391–401.

Mudgett, M.B. and Staskawicz, B.J. (1999) Characterization of the Pseudomonas syringae pv. tomato AvrRpt2 protein: demonstration of secretion and processing during bacterial pathogenesis. Mol Microbiol, 32, 927–941.

Nagar, R. and Schwessinger, B. (2018) DNA size selection (>3-4kb) and purification of DNA using an improved homemade SPRIbeads solution. Protocols.io, doi.org/10.17504/protocols.io.n7hdhj6.

Nakagawa, T., Suzuki, T., Murata, S., Nakamura, S., Hino, T., Maeo, K., Tabata, R., Kawai, T., Tanaka, K., Niwa, Y., Watanabe, Y., Nakamura, K., Kimura, T. and Ishiguro, S. (2007) Improved Gateway binary vectors: high-performance vectors for creation of fusion constructs in transgenic analysis of plants. Biosci Biotechnol Biochem, 71, 2095–2100.

Ntoukakis, V., Balmuth, A.L., Mucyn, T.S., Gutierrez, J.R., Jones, A.M. and Rathjen, J.P. (2013) The tomato Prf complex is a molecular trap for bacterial effectors based on Pto transphosphorylation. PLoS Pathog, 9, e1003123.

Oh, C.S. and Martin, G.B. (2011) Tomato 14-3-3 protein TFT7 interacts with a MAP kinase kinase to regulate immunity-associated programmed cell death mediated by diverse disease resistance proteins. J Biol Chem, 286, 14129–14136.

Oh, C.S., Pedley, K.F. and Martin, G.B. (2010) Tomato 14-3-3 protein 7 positively regulates immunity-associated programmed cell death by enhancing protein abundance and signaling ability of MAPKKKα. Plant Cell, 22, 260–272.

Pedley, K.F. and Martin, G.B. (2003) Molecular basis of Pto-mediated resistance to bacterial speck disease in tomato. Ann Rev Phytopathol, 41, 215–243.

Powell, A.F., Schmidt, M.H.-W., Courtney, L., Feder, A., Vogel, A., Xu, Y., Lyon, D., Dumschott, K., McHale, M., Bao, K., Duhan, A., Hallab, A., Denton, A.K., Mueller, L.A., Alekseh, S., Martin, C., Fernie, A.R., Martin, G.B., Fei, Z., Giovannoni, J.J., Strickler, S.R. and Usadel, B. (2020) A *Solanum lycopersicoides* reference genome provides resources for tomato researc. bioRxiv.

Prokchorchik, M., Choi, S., Chung, E.H., Won, K., Dangl, J.L. and Sohn, K.H. (2019) A host target of a bacterial cysteine protease virulence effector plays a key role in convergent evolution of plant innate immune system receptors. New Phytol, 225, 1327–1342.

Quast, C., Pruesse, E., Yilmaz, P., Gerken, J., Schweer, T., Yarza, P., Peplies, J. and Glockner, F.O. (2013) The SILVA ribosomal RNA gene database project: improved data processing and web-based tools. Nucleic Acids Res, 41, D590–596.

Roberts, R., Hind, S.R., Pedley, K.F., Diner, B.A., Szarzanowicz, M.J., Luciano-Rosario, D., Majhi, B.B., Popov, G., Sessa, G., Oh, C.S. and Martin, G.B. (2019) Mai1 protein acts between host recognition of pathogen effectors and mitogen-activated protein kinase signaling. Mol Plant Microbe Interact, 32, 1496–1507.

Rosli, H.G., Zheng, Y., Pombo, M.A., Zhong, S., Bombarely, A., Fei, Z., Collmer, A. and Martin, G.B. (2013) Transcriptomics-based screen for genes induced by flagellin and repressed by pathogen effectors identifies a cell wall-associated kinase involved in plant immunity. Genome Biol, 14, R139.

Saur, I.M., Conlan, B.F. and Rathjen, J.P. (2015) The N-terminal domain of the tomato immune protein Prf contains multiple homotypic and Pto kinase interaction sites. J Biol Chem, 290, 11258–11267.

Thapa, S.P. and Coaker, G. (2016) Genome sequences of two *Pseudomonas syringae pv. tomato* race 1 strains, isolated from tomato fields in California. Genome Announc, 4, e01671–01615.

Thapa, S.P., Miyao, E.M., David, M.R. and Coaker, G. (2015) Identification of QTLs controlling resistance to *Pseudomonas syringae* pv. tomato race 1 strains from the wild tomato, *Solanum habrochaites* LA1777. Theor Appl Genet, 128, 681–692.

Toruño, T.Y., Shen, M., Coaker, G. and Mackey, D. (2019) Regulated disorder: Posttranslational modifications control the RIN4 plant immune signaling hub. Mol Plant Microbe Interact, 32, 56–64.

Untergasser, A., Cutcutache, I., Koressaar, T., Ye, J., Faircloth, B.C., Remm, M. and Rozen, S.G. (2012) Primer3: new capabilities and interfaces. Nucleic Acids Res, 40, e115–e115.

Vogt, I., Wohner, T., Richter, K., Flachowsky, H., Sundin, G.W., Wensing, A., Savory, E.A., Geider, K., Day, B., Hanke, M.V. and Peil, A. (2013) Gene-for-gene relationship in the host-pathogen system Malus x robusta 5-*Erwinia amylovora*. New Phytol, 197, 1262–1275.

Whalen, M.C., Innes, R.W., Bent, A.F. and Staskawicz, B.J. (1991) Identification of *Pseudomonas syringae* pathogens of *Arabidopsis* and a bacterial locus determining avirulence on both Arabidopsis and soybean. Plant Cell, 3, 49–59.

Wick, R.R., Judd, L.M. and Holt, K.E. (2019) Performance of neural network basecalling tools for Oxford Nanopore sequencing. Genome Biol, 20, 129.

Wu, C.-H., Abd-El-Haliem, A., Bozkurt, T.O., Belhaj, K., Terauchi, R., Vossen, J.H. and Kamoun, S. (2017) NLR network mediates immunity to diverse plant pathogens. Proc Natl Acad Sci USA, 114, 8113–8118.

Wu, C.-H. and Kamoun, S. (2019) Tomato Prf requires NLR helpers NRC2 and NRC3 to confer resistance against the bacterial speck pathogen *Pseudomonas syringae* pv. tomato. bioRxiv, https://doi.org/10.1101/595744

Zimin, A.V., Marcais, G., Puiu, D., Roberts, M., Salzberg, S.L. and Yorke, J.A. (2013) The MaSuRCA genome assembler. Bioinformatics, 29, 2669–2677.

## References for Supplemental Information

Bej, A., Sahoo, B.R., Swain, B., M., B., Jayasankar, P. and Samanta, M. (2014) LRRsearch: An asynchronous server-based application for the prediction of leucine-rich repeat motifs and an integrative database of NOD-like receptors. Comp Biol Med, 53, 164–170.

Du, X., Miao, M., Ma, X., Liu, Y., Kuhl, J.C., Martin, G.B. and Xiao, F. (2012) Plant programmed cell death caused by an autoactive form of Prf is suppressed by co-expression of the Prf LRR domain. Mol Plant, 5, 1058–1067.

